# Systems Analysis of the 22q11.2 Microdeletion Syndrome Converges on a Mitochondrial Interactome Necessary for Synapse Function and Behavior

**DOI:** 10.1101/315143

**Authors:** Avanti Gokhale, Amanda A. H. Freeman, Cortnie Hartwig, Julia L. Bassell, Stephanie A. Zlatic, Christie Sapp, Trishna Vadlamudi, Farida Abudulai, Amanda Crocker, Erica Werner, Zhexing Wen, Gabriela M. Repetto, Joseph A. Gogos, Steven M. Claypool, Jennifer K. Forsyth, Carrie Bearden, Jill Gausier, David A. Lewis, Nicholas T. Seyfried, Victor Faundez

## Abstract

Neurodevelopmental disorders offer insight into synaptic mechanisms. To unbiasedly uncover these mechanisms, we studied the 22q11.2 syndrome, a recurrent copy number variant, which is the highest schizophrenia genetic risk factor. We quantified the proteomes of 22q11.2 mutant human fibroblasts and mouse brains carrying a 22q11.2-like defect, *Df(16)A+/-*. Molecular ontologies defined mitochondrial compartments and pathways as some of top ranked categories. In particular, we identified perturbations in the SLC25A1-SLC25A4 mitochondrial transporter interactome as associated with the 22q11.2 genetic defect. Expression of SLC25A1-SLC25A4 interactome components was affected in neuronal cells from schizophrenia patients. Furthermore, hemideficiency of the *Drosophila* SLC25A4 orthologue, dSLC25A4-*sesB*, affected synapse function and impaired sleep patterns in a neuronal-specific manner. These results identify a novel synaptic role of mitochondrial inner membrane solute transporters. We propose that mitochondria are among key organelles affected by genetic defects that increase the risk of neurodevelopmental disorders.

## Introduction

Single gene defects associated with neurodevelopmental disorders provide a fertile ground to uncover fundamental synaptic mechanism. For example, mutations in FMR1, MECP2, DISC1, or NRXN1 associate with diverse mental and/or behavioral disorders, including autism spectrum disorder and schizophrenia. Understanding of the molecular mechanisms linking these single gene defects with pathways that impinge on synapse function has been significantly advanced (Bena et al., 2013; Ishizuka et al., 2006; Santoro et al., 2012; Sztainberg and Zoghbi, 2016; Wen et al., 2014). This fact is founded on well-established experimental paradigms that identify and test causality between a single gene defect, its downstream molecular mechanisms, and phenotypes. In contrast, there are a great number of neurodevelopmental disorders associate with chromosomal microdeletions, which are hemizygous deletions containing multiple contiguous genes. Microdeletions have received great attention as they are the most penetrant and frequent genetic defects associated to neurodevelopmental disorders (Girirajan et al., 2011; Kirov, 2015; Malhotra and Sebat, 2012; Rutkowski et al., 2017; Sullivan et al., 2012). In comparison to monogenic disorders, the study of microdeletions is impeded by the lack of experimental paradigms that comprehensively capture contributions of all genes within the hemideletion to downstream molecular mechanisms and phenotypes (Iyer et al., 2018). Thus, the identity of molecular mechanisms downstream a whole microdeletion and their phenotypic impact in synapses remains elusive. Here we address this issue focusing on the 22q11.2 microdeletion syndrome.

The 22q11.2 microdeletion syndrome (OMIM #<u>192430</u>, #<u>188400</u>) (McDonald-McGinn et al., 2015) is the strongest and most prevalent genetic risk factor for schizophrenia increasing the overall risk of psychiatric pathology 20- to 25-fold as compared with the general population (Bassett and Chow, 2008; Bassett et al., 2000; Hodgkinson et al., 2001). Twenty five percent of 22q11.2 patients develop schizophrenia. In addition, the 22q11.2 microdeletion is the most common genetic defect found in sporadic cases of schizophrenia (Bassett and Chow, 2008; Bassett et al., 2003; Hoeffding et al., 2017; International Schizophrenia, 2008; Jonas et al., 2014; Karayiorgou et al., 2010; Marshall et al., 2017; Schneider et al., 2014). The strong association of mental and/or behavioral disorders with the 22q11.2 genetic defect makes this syndrome a robust model to test new experimental paradigms to identify molecular pathways and synaptic mechanisms downstream complex neurodevelopmental genetic defects.

We studied the most prevalent 22q11.2 microdeletion in humans, which encompasses three megabases. This microdeletion creates an haploinsufficiency of 46 protein coding genes and 17 regulatory small RNAs, thus opening the door for multiple pathways and organelles that could be affected downstream (Guna et al., 2015). We reasoned that top ranked molecular ontologies associated with the 22q11.2 genetic defect should enrich pathways and organelles implicated in mechanisms affecting synapse function and thus contribute to psychiatric phenotypes in humans. Using genealogical and integrated mass spectrometry based proteomics, we report the unbiased and statistically prioritized identification of pathways and organelles affected by the 22q11.2 microdeletion syndrome. Our comparative systems biology studies interrogated the proteome of fibroblasts from human pedigrees, genealogical proteomics, and the brain of a mouse model that genotypically and phenotypically mimics the 22q11.2 syndrome, the *Df(16)A+/-* deficiency (Karayiorgou et al., 2010). We conclude that the mitochondrion is a top ranked organelle affected in the 22q11.2 microdeletion syndrome.

## Results

### Genealogical and Comparative Proteomics Prioritize Mitochondrial Targets in 22q11.2 Microdeletions

We quantified proteome differences cosegregating with the 3 Mb microdeletion in 22q11.2 affected human fibroblasts and in brains from mice carrying a syntenic microdeletion in chromosome 16, *Df(16)A+/-*. We used human fibroblasts from pedigrees where one of the individuals was affected by 22q11.2 microdeletion syndrome and childhood psychosis, and compared affected subjects to their disease-free relatives. This strategy, termed genealogical proteomics, minimizes genetic variability between individuals and offers molecular insight into disease mechanisms despite limited subject number (Zlatic et al., 2018). We compared genealogical proteomic outcomes with *Df(16)A+/-* hippocampal and prefrontal cortex proteomes to identify universal mechanisms downstream of the 22q11.2 microdeletion.

We studied proteomes from the following families: one family where all members are disease-free (Fig. 1A), three families harboring one member affected by the 3 Mb 22q11.2 microdeletion (Fig. 1A, E, G), and two isolated 3 Mb 22q11.2 microdeletion patients (Fig. 1A and G). Proteomes were quantified with three mass spectrometry approaches: isobaric tandem mass tagging, (TMT, Fig.1B, F), triple SILAC (Fig. 1F, H), and label-free quantification (LFQ, Fig. 1F). The discriminatory power of genealogical proteomics was tested by comparing the cellular proteomes from 9 individuals within a single multiplexed TMT experiment. These nine individuals are organized in a disease-free family (Fig. 1A, subjects 11-16), a pedigree with one 22q11.2 affected subject (Fig. 1A, subjects 2-3), and an isolated 22q11.2 patient (Fig. 1A, subject 1). Hierarchical clustering of 4264 proteins quantified in all 9 subjects (Fig. 1B) segregated within a cluster all, but one, members of the unaffected family from unrelated subjects (Fig. 1C, subjects 11, 12, 14-16). This dataset contained the quantification of 10 out of the 46 proteins encoded within the 3 Mb 22q11.2 locus (Fig. 1D). Of these proteins SLC25A1, SEPT5, TXNRD2, COMT, RANBP1 and SNAP29 were predictably and significantly reduced by ~50% (Fig. 1D). Thus, genealogical proteomics discriminates genealogical relationships among a limited number of subjects and identifies expected protein expression levels in genes encoded within the 22q11.2 locus. Proteomic analysis of an independent 22q11.2 pedigree using three quantitative mass spectrometry approaches in independent experiments identified partially overlapping proteins whose expression was sensitive to the 22q11.2 microdeletion. However, these three datasets produced convergent and similarly ranked ontology terms (GO CC, see Canvas depiction in Fig. S1A). These results indicate that similar ontological inferences can be obtained from proteomic datasets produced by different quantitation methods, highlighting the rigor and reproducibility of our integrated proteomics approach.

**Fig. 1.**
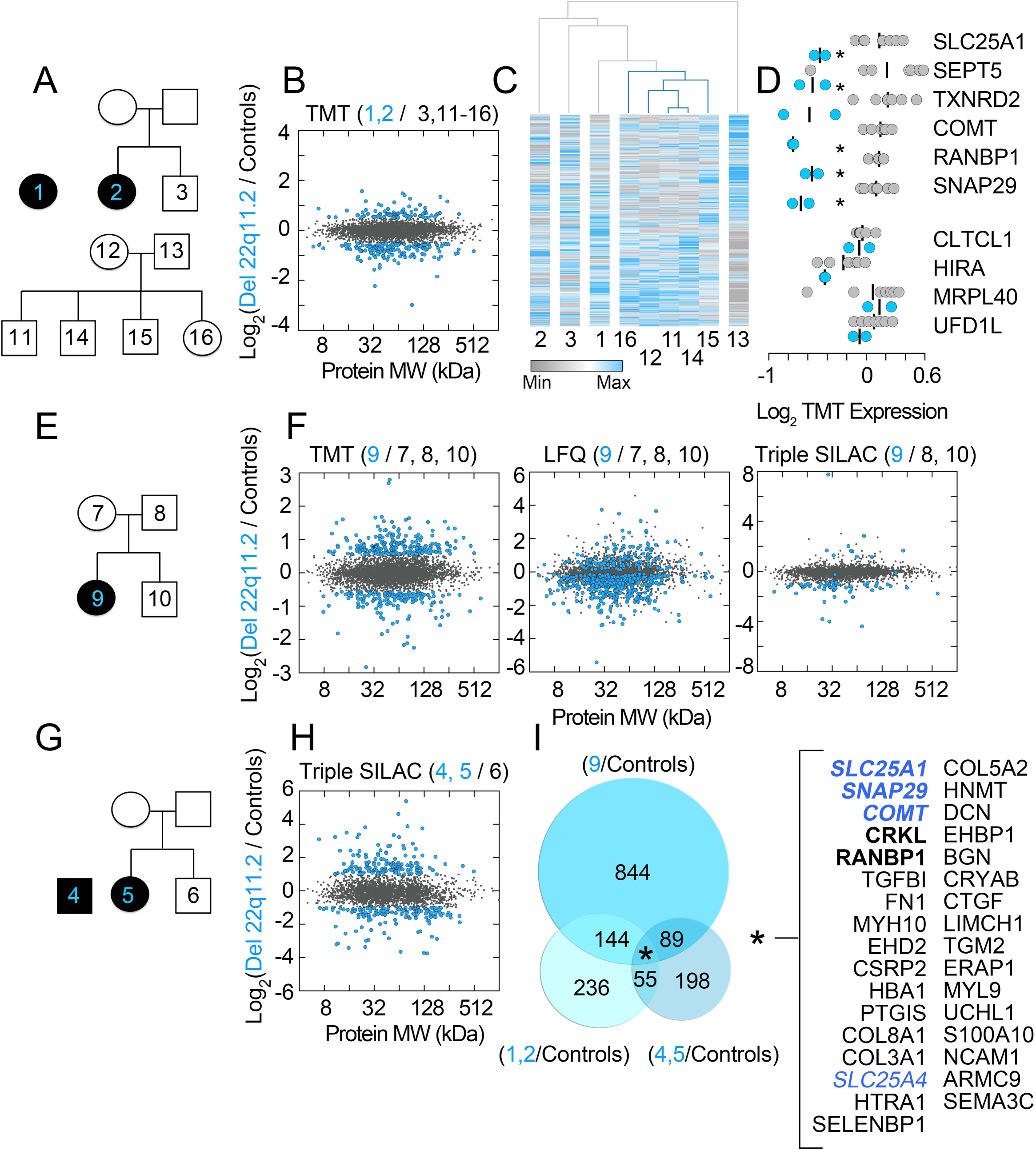
Genealogical Proteomics of 22q11.2 Pedigrees Fibroblasts Using Quantitative Mass Spectrometry. Human pedigrees of a control family (A) and families where one of the subjects is affected by 22q11.2 microdeletion syndrome and early childhood psychosis (blue numbers in A, E, G). Experimental design is designated at the upper left corner of dot plots (B, F, H). For example, (B) shows a TMT experiment where proteomes from probands 1 and 2 were compared against unaffected individuals 3, 11-16. C) Hierarchical clustering analysis of the proteome in subjects 1-3 and 11-16. Euclidian distance clustering of columns and rows (4264 TMT protein quantitations) shows segregation of related family members. D) Dot plot of proteins encoded within the 22q11.2 chromosomal segment quantitated in TMT experiment B. Asterisks denotes significant differences p= 0.04146 to p<0.0001 T-test. B, F and H depict all mass spectrometry quantifications where the color code denotes individuals being compared (blue symbols are proteins whose expression is changed, grey symbols are unaffected proteins). Significant protein expression changes for: TMT and SILAC were consider to be >2 or <0.5 whereas in label free quantification (LFQ) a -log(p) value threshold of 1.3 was used. I) Venn diagram summarizes proteins with significant expression changes in B, F and H. Asterisk denotes proteins whose expression changed in all patients. Bold font depicts proteins encoded within the 22q11.2 segment. Blue color fonts are proteins contained in the human Mitocarta 2.0 dataset. Individual MS/MS data can be found in Table S1.

Genealogical proteomics of the three 22q11.2 pedigrees (Fig. 1A-E-G) collectively identified 1500 proteins whose expression was altered in 22q11.2 microdeletion cells (Fig. 1I). Of these proteins, only 18 polypeptides were common to all of the 22q11.2 affected individuals (Fig. 1I), including five polypeptides contained in the 22q11.2 locus and 13 polypeptides previously not implicated in 22q11.2 syndrome (Fig. 1I). Independent gene ontology analysis of each one of these three pedigree datasets converged on partially overlapping gene ontology categories (Fig. S1B and C). We inferred ontological categories with the 1500 proteins whose expression cosegregated with the 22q11.2 microdeletion, hereafter referred as the 22q11.2 proteome. We used three bioinformatic algorithms which produced similarly ranked ontological categories. We queried the gene ontology cellular component (GO CC), REACTOME, and KEGG pathways simultaneously with the ClueGo algorithm to discern statistically ranked organelles, pathways, and associated pathologies downstream of the 22q11.2 microdeletion (Bindea et al., 2009). The top ontology categories/pathways were all related to mitochondrial compartments (Fig. 2A and Table S2, group q value 1.05E-38), as well as diseases where mitochondria are implicated in pathogenesis such as Parkinson’s and Huntington’s disease (Fig. 2A and Table S2, group q value 3.93E-37) (Lin and Beal, 2006). Additionally, the 22q11.2 proteome was enriched in extracellular matrix, lysosome, and actin cytoskeleton components and pathways (Fig. 2A and Table S2, group q values 3.23E-23, 6.61E-18, and 7.93E-16, respectively). We confirmed these bioinformatic results with the ENRICHR engine to interrogate the KEGG, OMIM, and GO CC databases (Chen et al., 2013). Mitochondrial compartments and pathways, Parkinson’s, Huntington’s and other diseases where mitochondria are affected were enriched in the 22q11.2 proteomic dataset (Fig. 2B, q values 3.3E-36, 1.2E-21, and 5.1 E-12; respectively).

**Fig. 2.**
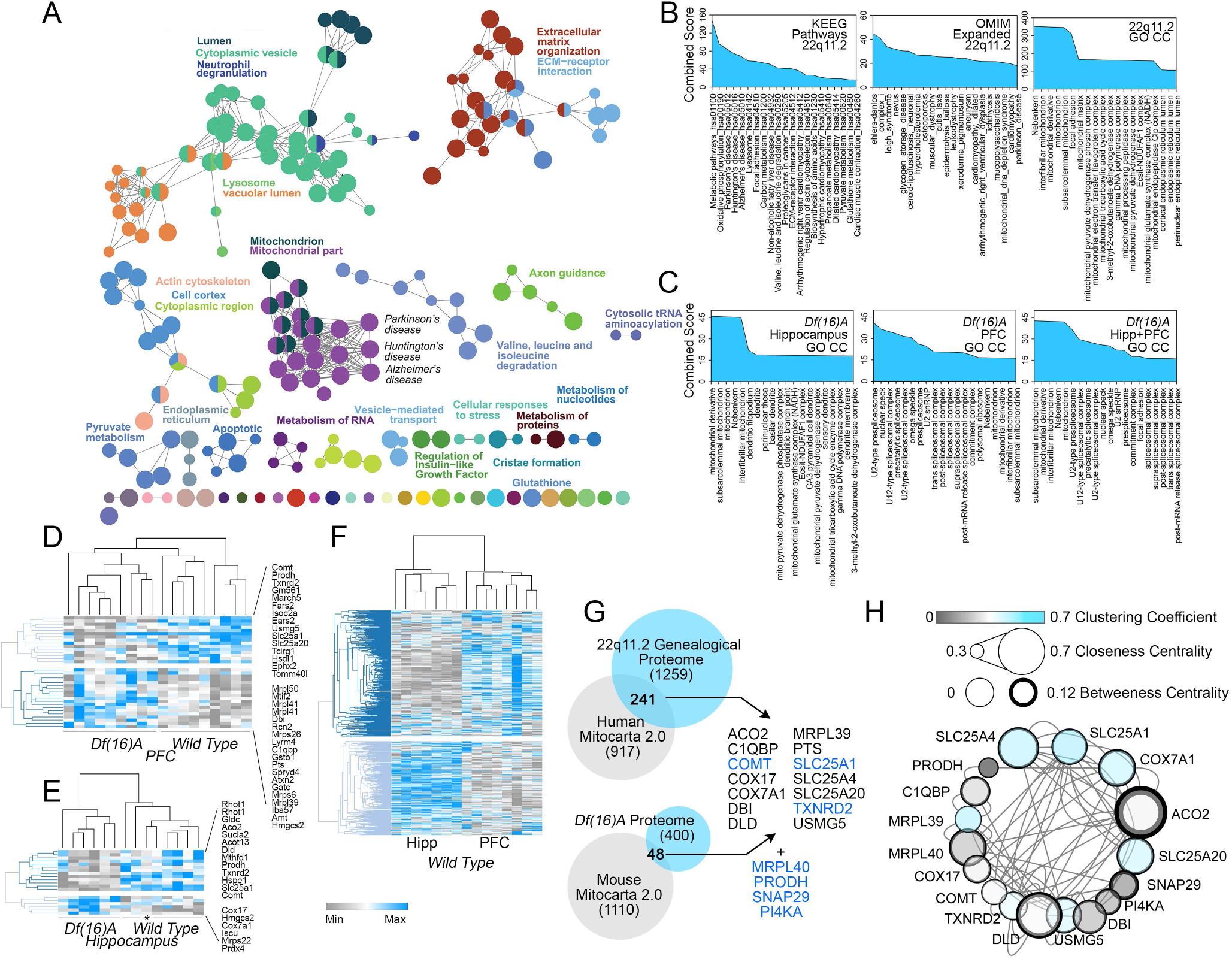
Comparative Bioinformatic Analysis of the 22q11.2 and the *Df(16)A+/-* Mouse Brain Proteomes. A) The 22q11.2 Proteome was analyzed with the engine ClueGo integrating the Cellular Component gene ontology GO:CC, REACTOME and KEGG databases. Functionally grouped network was built with terms as nodes and edges based on their term-term similarity statistics. The node size represents the term enrichment significance (p < 0.015 Bonferroni corrected). B) ENRICHR analysis of the 22q11.2 Proteome querying GO CC, KEEG and OMIM databases. C) ENRICHR analysis of the *Df(16)A+/-* mouse hippocampus and prefrontal cortex (PFC) proteomes as in B. Mouse brain proteomes were quantified using TMT mass spectrometry (see table S3, n=6 mutant and 5 control mice). D-F) Differences in the mitoproteomes of wild type and *Df(16)A+/-* mitoproteomes. Hierarchical clustering analysis of the *Df(16)A+/-* mouse hippocampus and prefrontal cortex mitochondrial proteome hits (D-E) is compared to the wild type mitoproteomes quantified in hippocampus (Hipp) and prefrontal cortex (PFC). Kendall’s tau distance clustering of columns and rows. G) Venn diagrams present overlapping protein hits between the 22q11.2 and *Df(16)A+/-* proteome with the human and mouse Mitocarta 2.0 datasets. Listed proteins correspond to mitochondrial proteins whose expression is sensitive to the microdeletion in human and mouse (upper two columns). Lower column and blue font proteins are encoded in the 22q11.2 chromosomal segment. H) SLC25A1 and SLC25A4 are high connectivity nodes in a discrete 22q11.2 and *Df(16)A+/-* mitoproteome interactome. In silico interactome of protein hits listed in G. Interactome was analyzed with graph theory to determine high connectivity nodes predictive of essential genes. Additional bioinformatic data and MS/MS data can be found in Tables S2-S3.

We examined ontology terms inferred from a brain proteome sensitive to the syntenic *Df(16)A+/-* deficiency in mice, hereafter referred as the *Df(16)A+/-* brain proteome (Fig. 2C). We reasoned that overlapping ontological categories between the 22q11.2 proteome and the *Df(16)A+/-* brain proteome would point to robust and universal mechanisms downstream of the 22q11.2 microdeletion. We profiled by TMT the hippocampus and prefrontal cortex proteomes of control and *Df(16)A+/-* mouse brains. We quantified 6419 proteins and identified 110 hippocampal and 365 prefrontal cortex proteins whose expression was sensitive to the *Df(16)A+/-* microdeletion. ENRICHR bioinformatic analysis indicated that mitochondrial terms were top ranked in the *Df(16)A+/-* hippocampus proteome (Fig. 2C, q value 0.0018 and combined score of 45.77). In contrast, the spliceosome ranked first in the *Df(16)A+/-* prefrontal cortex proteome (Fig. 2C, q value 5.01E-06 and combined score of 41.16) with mitochondrial ontological categories scoring in the sixteenth place (Fig. 2C, p value 0.012 and combined score of 16.52). These ranking differences among ontological hits were due to different mitochondrial polypeptides being affected by the *Df(16)A+/-* microdeletion in the hippocampus and prefrontal cortex mitoproteomes (Fig. 2D-E). We attribute these differences to distinctive mitoproteome stoichiometries that distinguish the hippocampus and prefrontal cortex in control mouse brain (Fig. 2F). These regional mouse brain mitoproteome differences were also observed in flies were distinct neurons of the *Drosophila* mushroom body, the fly hippocampus equivalent (Campbell and Turner, 2010), can be segregated away just based on stoichiometric differences in the transcriptome encoding the fly mitoproteome (Fig. S2) (Chen et al., 2015; Crocker et al., 2016). We conclude that the proteomes sensitive to either the 22q11.2 or the *Df(16)A+/-* hemideficiencies enrich components of the mitoproteome in a brain region specific manner.

### Identification and Prioritization of Key Mitochondrial Proteins within the 22q11.2 Proteome

We used the Mitocarta 2.0 mitoproteome dataset as a reference to identify mitochondrial proteins among the 22q11.2 and *Df(16)A+/-* proteomes (Calvo et al., 2016; Pagliarini et al., 2008). We identified 241 mitochondrial proteins sensitive to the 22q11.2 microdeletion and 48 mitochondrial proteins sensitive to the *Df(16)A+/-* deficiency (Fig. 2G). Expression of fourteen mitochondrial proteins was affected either in all human pedigrees (Fig 1I) or simultaneously in human and mouse cells with the microdeletion (Fig. 2G). We merged these 14 mitochondrial proteins with four additional proteins encoded within the 22q11.2 chromosomal segment which are also part of the Mitocarta 2.0 datasets. A network of protein-protein interactions constrained to these 18 polypeptides was subjected to graph theory analysis to unbiasedly determine node relevance within this network (Fig. 2G, blue font represents 22q11.2 encoded proteins, and Fig. S3). We used clustering, closeness centrality, and betweenness centrality coefficients to measure node relevance (del Rio et al., 2009). The gene products with the highest relevance scores within this interactome were SLC25A1 and SLC25A4. SLC25A1 and SLC25A4 are encoded in the 22q11.2 and 4q35.1 cytogenetic bands. These two transporters participate in central inner mitochondrial solute transport mechanisms and are widely expressed in multiple tissues (Palmieri and Monne, 2016; Taylor, 2017). Thus, we selected these two inner mitochondrial transporters as candidate genes whose disruption would maximize network perturbation.

We confirmed that SLC25A1 and SLC25A4 expression were altered in 22q11.2 fibroblasts as compared to non-affected family members. Both transporters were decreased at least by 50% in 22q11.2 affected fibroblasts as compared to unaffected family members (Fig. 3A, compare lanes 1 and 2, 3-4 and 5, quantified in Fig. 3B). We hypothesized that co-expression changes observed in microdeletion patient cells may be the result of biochemical/metabolic interactions between SLC25A1 and SLC25A4. We used two approaches to address this question. First, we tested whether SLC25A1 and SLC25A4 influenced each other’s expression, a common occurrence in proteins that physically interact or belong to a pathway (Wu et al., 2013). We used cells where SLC25A1 or SLC25A4 expression was abrogated by CRISPR-Cas9 genome editing. Cells lacking SLC25A1 significantly increased the expression of SLC25A4 ~1.5-2-fold, while SLC25A4-null cells upregulated SLC25A1 3.6 times demonstrating a genetic interaction between these two transporters (Fig. 3C-D). Second, we performed immunomagnetic isolation of SLC25A1 from detergent soluble extracts from wild type and either SLC25A1 or SLC25A4 mutant cells. An SLC25A1 antibody robustly immunoprecipitated a SLC25A1 immunoreactive band absent in SLC25A1 null cells (Fig. 3E compare lanes 3-4). This SLC25A1 antibody also co-immunoprecipitated SLC25A4 from wild type cell extracts but not from SLC25A4 null cells (Fig. 3F compare lanes 3-4). We determined co-precipitation selectivity by blotting for transferrin receptor, a transmembrane protein absent from Mitocarta 2.0 (Fig. 3F, TFRC) (Calvo et al., 2016; Pagliarini et al., 2008). Reverse immunomagnetic isolations with FLAG-tagged SLC25A4 and its paralog SLC25A5 recovered endogenous SLC25A1 (Fig. 3G, lanes 2-5). SLC25A1 co-isolation with tagged SLC25A4 and 5 was prevented by FLAG peptide competition (Fig. 3G, lanes 3-6). Collectively, these findings demonstrate that SLC25A1 and SLC25A4 genetically and biochemically interact.

**Fig. 3.**
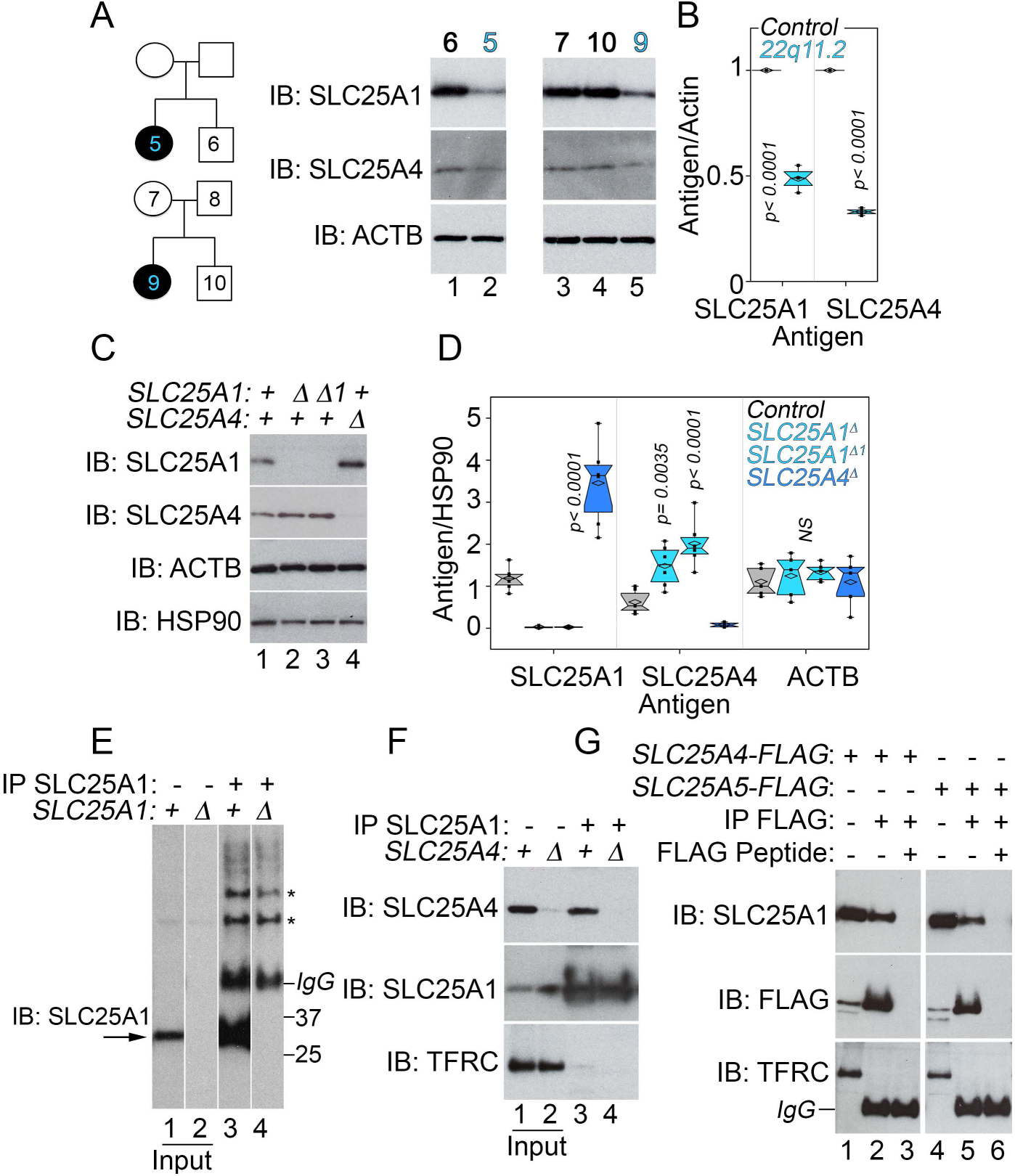
SLC25A1 and SLC25A4 Expression is Affected by the 22q11.2 Microdeletion Cell and these transporters Biochemically and Genetically Interact. A) Human pedigrees of families affected by 22q11.2 microdeletion syndrome. Immunoblots of total cellular lysates from fibroblasts obtained from individuals in pedigrees. B) Quantitation of results shown in A. p values One way ANOVA followed by Dunnett’s multiple comparisons, n=3. C) SLC25A1 and SLC25A4 expression changes in cells carrying null mutations (∆) in *SLC25A1* or *SLC25A4* clonal cell lines. Detergent soluble cell extracts were blotted with indicated antibodies. Actin (ACTB) and HSP90 were used as controls. D) Depicts quantitation of expression levels as compared to wild type cells. p values One way ANOVA followed by Dunnett’s multiple comparisons, n=5. E) SLC25A1 antibody precipitates an SLC25A1 immunoreactive band (lane 3) absent from SLC25A1 null cells (lane 4). Asterisks denote non-specific bands recognized by the antibody. F) SLC25A1 antibody precipitates an SLC25A4 immunoreactive band (lane 3) absent from SLC25A4 null cells (lane 4). G) FLAG tagged SLC25A4 or SLC25A5 precipitate SLC25A1 (lames 2 and 5). Lanes 1 and 3 correspond to inputs. Lanes 4 and 6 correspond to immunoprecipitation where an excess FLAG peptide was used for outcompetition. F-G) Transferrin receptor (TFRC) was used as a control for non-specific membrane protein precipitation.

### Expression of SLC25A Family of Mitochondrial Transporters is Altered in 22q11.2 Fibroblasts and Schizophrenia Patient Neurons

We created a comprehensive *ab initio* SLC25A1-SLC24A4 interactome using as building blocks a SLC25A4-focused interactome plus all SLC25A1 and SLC25A4 interactions curated from seven proteome wide physical interaction datasets (Fig. 4A and S3) (Floyd et al., 2016; Havugimana et al., 2012; Hein et al., 2015; Huttlin et al., 2017; Huttlin et al., 2015; Lu et al., 2017; Wan et al., 2015). The *ab initio* SLC25A1-SLC25A4 interactome contained 106 nodes encompassing mitochondrial respiratory chain components and twelve SLC25A transporter family members (Fig. 4A and S3). The SLC25A1 and SLC25A4 nodes maintained their relative relevance within the *ab initio* network, as ascertained by SLC25A1 and SLC25A4 centrality coefficients (Figs. 4A and S3). Forty five of the 106 *ab initio* SLC25A1-SLC25A4 interactome nodes were represented in the human 22q11.2 proteome indicating a convergence of the 22q11.2 proteome mitochondrial hits and the *ab initio* network (Figs. 4A and S3, grey nodes).

**Fig. 4.**
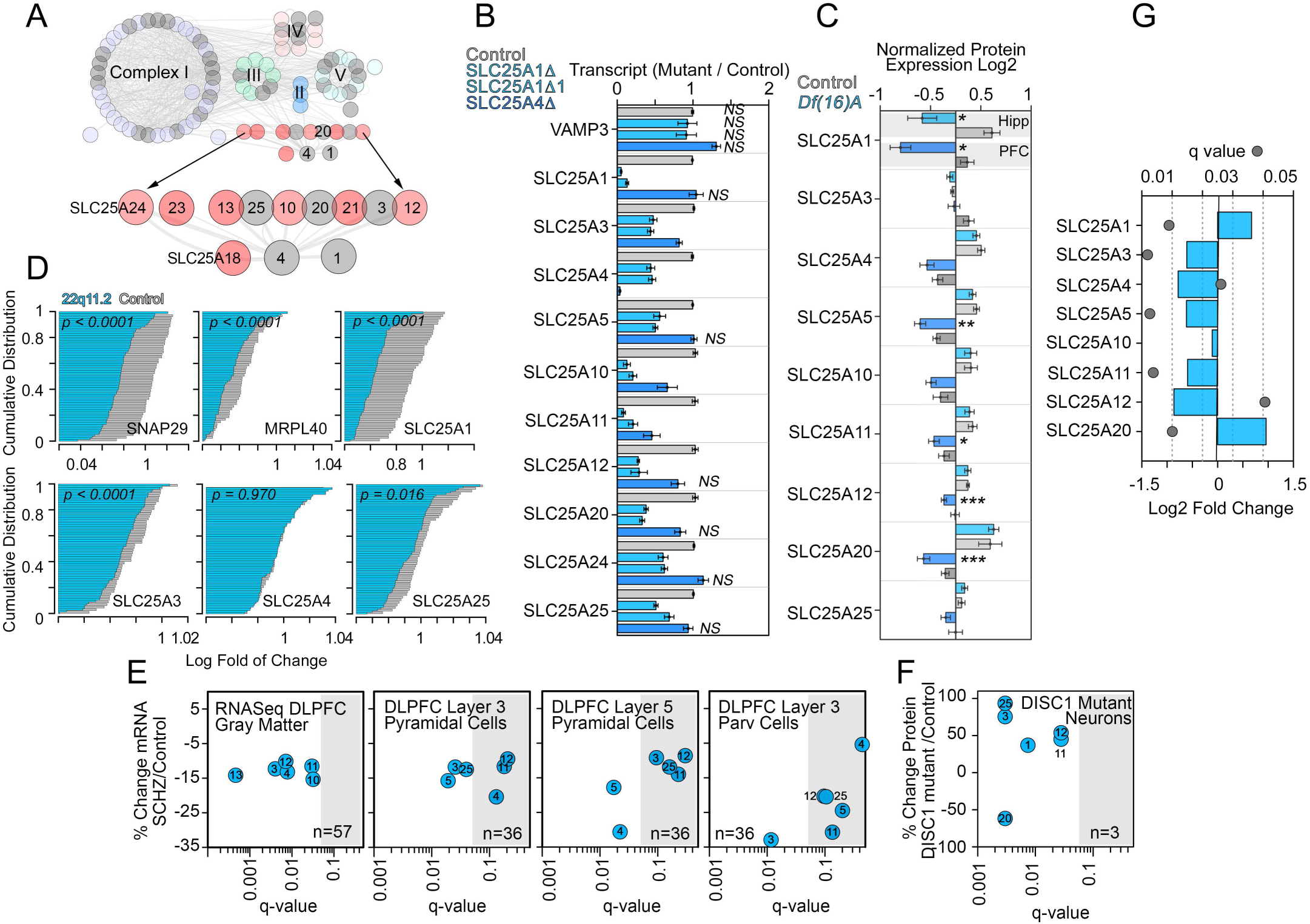
Expression of Components of the SLC25A1-SLC25A4 Interactome is Affected in Neurodevelopmental Disorders. A) Comprehensive *in silico* interactome of the SLC25A1 and SLC25A4 mitochondrial transporters. Complexes I to V of the respiratory chain as well as SLC25A transporter family members are color coded. All nodes colored gray represents hits in the 22q11.2 proteome. Additional details can be found in figure S3A. B) Expression of SLC25A transporter family member transcripts is altered in SLC25A1 or SLC25A4 null cells. Transcript quantification by qRT-PCR is expressed as ratio to vimentin mRNA. VAMP3 was used as control. n=4 One Way ANOVA followed by Fisher’s Least Significant Difference Comparison. All non-significant comparisons are marked (NS). C) Expression of SLC25A transporter family member polypeptides is altered in *Df(16)A+/-* mouse hippocampus (Hipp) or prefrontal cortex (PFC). SLC25A transporters were quantitated by TMT mass spectrometry. n= 6 mutant and 5 control mice. One Way ANOVA followed by Fisher’s Least Significant Difference Comparison. Asterisk marks p≤0.0001. ^**^ p=0.0098, ^***^ p≤0.028. D) Expression of SLC25A family member mRNAs is reduced in whole blood from unaffected and 22q11.2 patients. Probability plots of mRNA quantified by microarray on 50 unaffected (grey) and 77 22q11.2 patients (blue). SNAP29, MRPL40 and SLC25A1 reside in the 22q11.2 microdeletion locus and were used as controls to determine the range of expression change attributable to the microdeletion. SLC25A3 and SLC25A25 expression is modified within this range. P values were calculated using Kolmogorov–Smirnov test. E) mRNA expression of SLC25A transporters in gray matter or single cells isolated from unaffected and schizophrenia cases. Gray matter mRNA quantitations were performed by RNAseq while single cell mRNA quantitations were performed by microarray in dorsolateral prefrontal cortex (DLPFC) samples. F) Proteomic quantitation of SLC25A transporters in iPSC-derived cortical neurons from DISC-1 mutant patient and isogenic controls. E-F) SLC25A transporter family members SLC25An where n correspond to the number on blue circle. Grey box denotes non-significant changes in expression after multiple corrections. G) mRNA expression of SLC25A transporter family members is altered in schizophrenia brains. Meta-analysis data obtained from Gandal et al. (Gandal et al., 2018)

We selected the SLC25A family member transporters to test the reliability of the *ab initio* network (Fig. 4A). We asked whether members of the SLC25A transporter family genetically interacted as inferred from the *ab initio* SLC25A1-SLC25A4 interactome. We first investigated if mRNA levels of SLC25A transporter family members were altered in SLC25A1- or SLC25A4-null cells. We measured transcripts of 10 of the 12 *ab initio* network SLC25A transporters in both SLC25A1 and SLC25A4 knock out cells by qRT-PCR. SLC25A1 null cells significantly altered the expression of 9 of the 10 measured SLC25A transporters while SLC25A4 null cells affected three transporters (Fig. 4C). These changes in transcript content were selective as evidenced by unaltered levels of the housekeeping genes VAMP3 and VIM (Fig. 4B). We further analyzed if these SLC25A network transporters were affected in other 22q11.2 and syntenic microdeletion tissues. Quantitative mass spectrometry of SLC25A family transporters showed an anticipated decrease of ~50% in SLC25A1 in prefrontal cortex and hippocampus of *Df(16)A+/-* mice (Fig. 4C). Additionally, expression of five out of nine SLC25A family transporters was decreased in prefrontal cortex (Fig. 4C). We extended these observations to lymphoblasts from 77 22q11.2 microdeletion patients and compared mRNA levels to 50 unaffected subjects. Expression of genes within the 22q11.2 locus significantly reduced their expression (Fig. 4D. SNAP29, MRPL40 and SLC25A1). Messenger RNA levels of three out of 10 SLC25A family transporters were significantly reduced in 22q11.2 cells (Fig. 4D). We conclude that components of the *ab initio* SLC25A1-SLC25A4 interactome are affected in tissues from human and mouse carrying 22q11.2 and syntenic microdeletions, respectively. These findings validate the *ab initio* SLC25A1-SLC25A4 interactome for studies of neurodevelopmental and synaptic mechanisms.

We hypothesized that if components of the *ab initio* SLC25A1-SLC25A4 interactome were to participate in the pathogenesis or phenotypic expression of 22q11.2 neuropsychiatric nosology, then neurons from patients affected by schizophrenia or neurons from patients with mutations in the schizophrenia risk gene DISC1 would alter the expression of SLC25A transporters. Expression of SLC25A transcripts was measured by RNAseq in dorsolateral prefrontal cortex gray matter of 57 of age- and sex-matched pairs of unaffected comparison and schizophrenia subjects (Fig. 4E). Of the six SLC25A family transporters with detectable levels of expression, all had mean mRNA levels that were lower in subjects with schizophrenia (Fig. 4E, q<0.05 SLC25A3, SLC25A4, SLC25A11, and SLC25A12). Next, pools of layer 3 and layer 5 pyramidal cells and layer 3 parvalbumin cells were individually collected via laser capture microdissection in a subset of subjects (N=36 pairs), and SLC25A transcripts were measured via microarray (Arion et al., 2015; Enwright Iii et al., 2017). Expression of each of the six SLC25A transcripts that were detectable by microarray was lower in all three cell types, though not all met statistical significance (Fig. 4F). The most affected transporter in schizophrenia layer 5 pyramidal neurons was SLC25A4 (Fig. 4E, 30.6% reduction, q value = 0.0222) while in parvalbunim neurons it was SLC25A3 (Fig. 4E, 32.9% reduction, q value = 0.0116). Changes in the expression of SLC25A family transporters was also observed in iPSC-derived human prefrontal neurons carrying a frameshift mutation of DISC1 as compared to isogenic controls generated by editing of the DISC1 gene defect (Wen et al., 2014). Proteomics identified significant changes in the expression of six out of 12 *ab initio* network SLC25A transporters with the most pronounced effects on SLC25A3, SLC25A20, and SLC25A25 (Fig. 4F, q values <0.003). These findings are also reproduced in a recent and comprehensive metanalysis of mRNA expression changes in 159 cortical schizophrenia patient samples compared to 293 unaffected subjects (Fig. 4G) (Gandal et al., 2018). These results indicate that components of the *ab initio* SLC25A1-SLC25A4 interactome are altered in tissue samples from patients affected by neurodevelopmental disorders sharing phenotypes with the 22q11.2 syndrome. Collectively, these findings demonstrate that SLC25A1 and SLC25A4 are principal nodes within a mitochondrial interactome. Our results suggest that these mitochondrial transporters and their interactome may participate in mechanisms necessary for synapse function and behavior.

### SLC25A1 and SLC25A4 genetic and functional interactions are required for normal synapse development, function, and behavior

To test the role of SLC25A mitochondrial transporters on neuronal function and behavior, we selected *Drosophila* because of genetic tools that allow precise control of gene expression in a developmental-, cell-, and tissue-restricted fashion. In particular, we focused on studying developmental synaptic and adult sleep behavioral phenotypes caused by changes in the expression of SLC25A1 and SLC25A4 in *Drosophila*. SLC25A1 and SLC25A4 orthologues are encoded by the gene *scheggia* (*sea*, CG6782, dSLC25A1) and *stress-sensitive B* genes (*sesB*, CG16944, dSLC25A4), respectively. We examined functional, electrophysiological phenotypes using the third instar *Drosophila* neuromuscular junction synapse, which is a reliable model to assess synaptic developmental phenotypes associated with neurodevelopmental gene defects (Frank et al., 2013). We controlled the expression of dSLC25A1-sea with UAS-RNAi reagents. dSLC25A4-sesB expression was modified with UAS-RNAi as well as two genomic alleles of *sesB, sesB^org^* and *sesB^9Ed-1^*. *sesB^org^* is a thermosensitive allele that does not affect synaptic mitochondrial content at permissive temperatures; however, the ADP-ATP transport activity decreases by 60%, offering a functional hemideficiency model (Rikhy et al., 2003). In contrast, *sesB^9Ed-1^* is a strong lethal null allele that is viable as single copy deficiency (Zhang et al., 1999). We used *sesB^org^* at the permissive temperature and *sesB^9Ed-1^/+* to mimic the partial reduction in the expression of SLC25A family members observed in samples from patients diagnosed with schizophrenia (Fig. 4). We drove the expression of *UAS-sea* and *UAS-sesB* RNAi transgenes using neuronal specific *elav^c155^-Gal4* and *VGlut-Gal4* transgenic drivers and analyzed the morphology of the larval neuromuscular junction (Fig. 5). Reduced neuronal expression of sea or sesB increased the number of boutons and branches per synapse (Fig. 5). The genomic alleles *sesB^org^*, at permissive temperature, and *sesB^9Ed-1^*/+ phenocopied neuronal specific RNAi-mediated down-regulation of dSLC25A4-sesB (Fig. 5) demonstrating that partial compromise of dSLC25A4-sesB expression alters synapse morphology. Despite these morphological changes in *sesB* mutant synapses, neurotransmission was not altered (Fig. 6). Evoked neurotransmission, measured as evoked excitatory junction potential amplitude (EJP) (Fig. 6A and C), and spontaneous miniature excitatory junction potential (mEJP), measured by the frequency and amplitude (Fig. 6B and D), remained unaffected in all *sesB* genotypes tested.

**Fig. 5.**
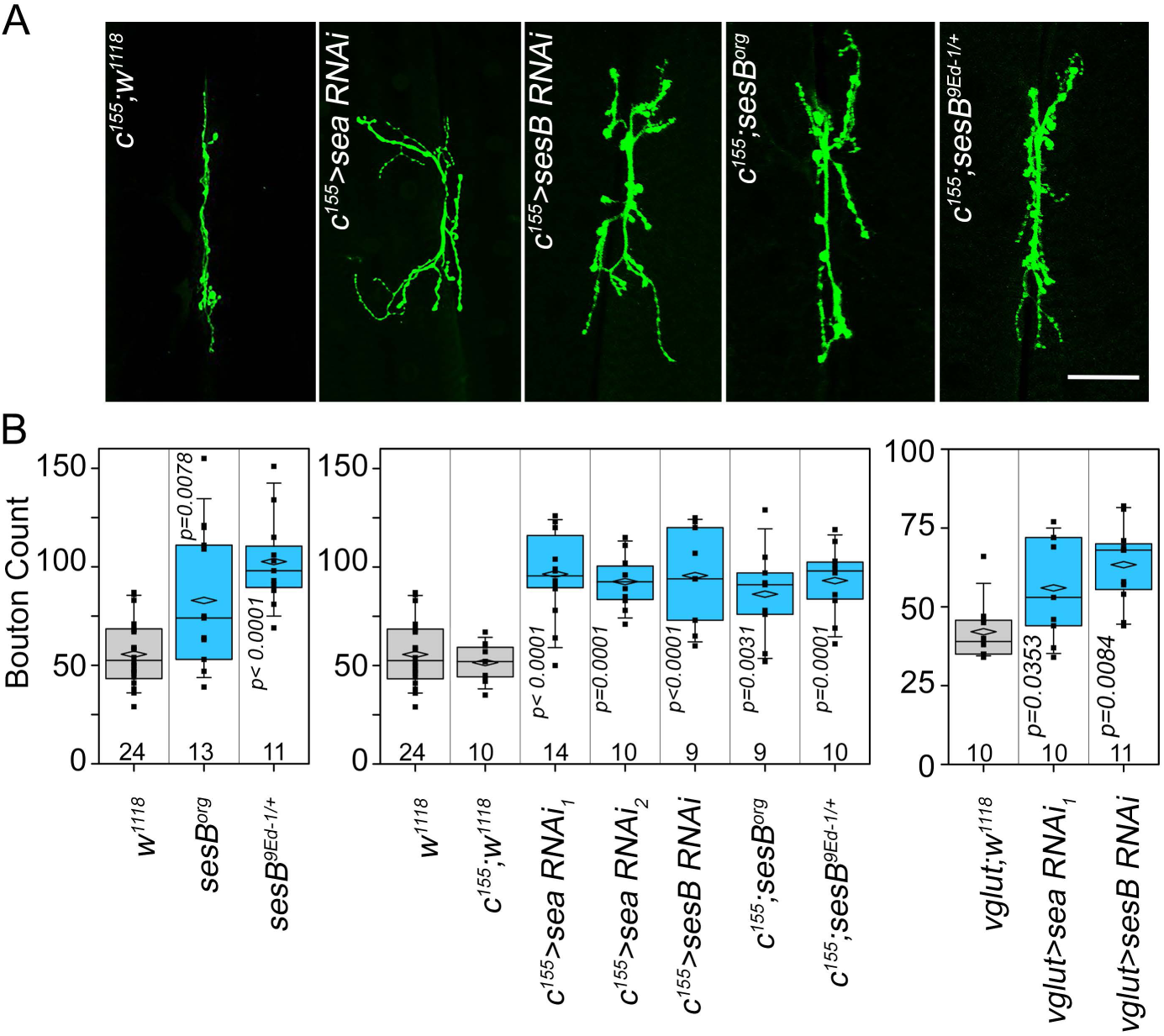
Reduced Expression of *Drosophila* dSLC25A1-dSLC25A4 Alters Synapse Morphology. A and B) Muscle VI-VII third instar neuromuscular junctions were stained with antibodies against the neuronal marker HRP. Expression of SLC25A1 (*sea*, *scheggia*) was downregulated with two RNAi transgenes. dSLC24A4 expression was reduced with a RNAi transgene or two genomic alleles (*sesB^org^* and *sesB^9Ed-1/+^*). Neuronal specific expression of RNAi regents was driven by the *elav^c155^*-GAL4 driver (*c155*). w1118 or w1118; *elav^c155^*-GAL4 animals were used as controls B) Shows quantitation of bouton counts per synapse. Counts were performed blind to the animal genotype. All comparisons in B were performed with One-Way ANOVA followed by Bonferroni’s multiple comparison.

**Fig. 6.**
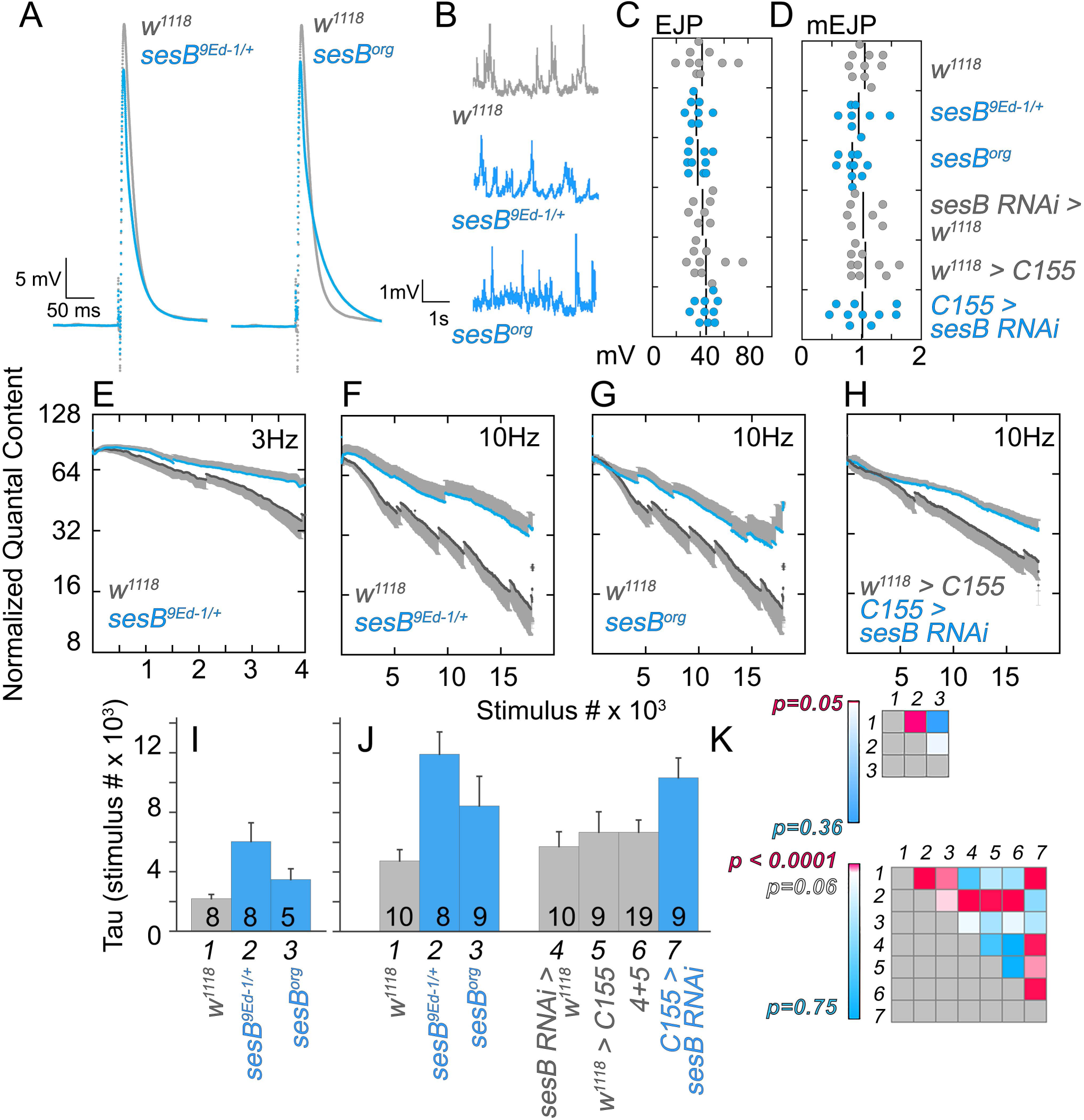
Hemideficiency of the *Drosophila* SLC25A4 Orthologue, *sesB,* Impairs Synapse Function. Muscle VI-VII third instar neuromuscular junctions from w1118 control (gray traces) and *sesB* mutants (blue traces) were analyzed for evoked (EJP in A) and spontaneous neurotransmission (mEJP in B). C and D present amplitudes as dot plots with each dot corresponding to one animal. Lines depicts the mean of the sample. E) Neuromuscular junctions stimulated at low frequency (3 Hz, E) and high frequency (10 Hz, F-H) in the presence of 1μM bafilomycin A1 to assess recycling and reserve pools of synaptic vesicles. Graphs E-H show control animals as black symbols (w1118, E-G; w1118>C155, H), blue symbols show *sesB* mutants (F-G), and neuronal specific sesB RNAi (H, C155>sesB RNAi). Average±SEM. I-J) shows quantitation of graphs E-H as time (measured as stimulus number) to 50% depletion (Tau) compared to response at time/stimulus 0. I) correspond to synapses stimulated at 3Hz, recycling pool of vesicles while J) shows results for synapses stimulated at 10Hz, reserve pool of vesicles. Number of animal is shown in the bar bottom. Average ± SEM. K) Tau statistical differences among genotypes at 3Hz (upper panel) and 10Hz (lower panel) represented as heat maps. Italic numbers depict genotypes in I) and J). All comparisons in I and J were performed with One-Way ANOVA followed by Fishers’s multiple comparison.

*Drosophila* genetic defects in genes associated with neurodevelopmental risk perturb neurotransmission at the neuromuscular junction at high frequency while keeping EJPs and mEJPs unaltered (Chen et al., 2017; Mullin et al., 2015). A similar synaptic phenotype is observed in a fly mutant affecting synaptic mitochondrial fission (Verstreken et al., 2005). Therefore, we examined and compared neurotransmission elicited at 3Hz and at high frequency (10Hz) on wild type and *sesB* deficient synapses (Fig. 6E-K). We incubated neuromuscular junctions in the presence of bafilomycin A1, a vacuolar ATPase inhibitor, to prevent neurotransmitter vesicle reloading after a round of vesicle fusion (Delgado et al., 2000; Kim et al., 2009; Mullin et al., 2015). This strategy leads to synapse fatigue in wild type larvae due to synaptic vesicle depletion (Fig. 6E-H, gray curves). Synaptic transmission at low frequency stimulation was normal in *sesB* deficient synapses (Fig. 6E, I, K). However, neurotransmission at high frequency was consistently increased in all *sesB* alleles as expressed by synaptic resilience to fatigue (10Hz, Fig. 6F-G, J, K). These effects were due to changes in the expression of sesB in neurons and not the muscle, as demonstrated by down-regulation of sesB with neuronal-specific RNAi (Fig. 6H, J, K). These results demonstrate that partial reduction in the expression of SLC25A4 modifies steady state morphology and increases synapse resilience to the increased functional demand of a developing synapse.

To assess the behavioral consequences of partial loss of function in components of the SLC25A1-SLC25A4 interactome in adults, we analyzed sleep patterns in wild type animals and mutants carrying *sesB^org^*, *sesB^9Ed-1^*, and *sesB* RNAi (Fig. 7). We also contrasted *sesB*-dependent phenotypes with sleep phenotypes induced by down-regulation of dSLC25A1-sea (Fig. S4). We chose sleep as a non-invasive and high throughput analysis of an adult behavior which is entrained by environmental cues (Freeman et al., 2012; Hendricks et al., 2000). Moreover, sleep alterations are frequent in neurodevelopmental disorders (Chouinard et al., 2004; Kamath et al., 2015; Krakowiak et al., 2008; Monti and Monti, 2004; Petrovsky et al., 2014). We measured locomotor activity using the *Drosophila* Activity Monitoring (DAM) system to quantify episodes of activity and sleep in a 12:12 hour light:dark cycle. Wild type Canton S animals demonstrated the highest density of sleep activity during the dark period (Fig. 7A-B and Fig. S4, zeitgeber times ZT12 to 24) with an increased number of sleep-wake transitions at the beginning and end of the light cycle (Fig. 7A-B and Fig. S4, zeitgeber times ZT1 and 12). This pattern was disrupted in *sesB^org^* and *sesB^9Ed-1^* animals, which exhibit increased sleep-wake transitions throughout the 24-hour period (Fig. 7A-B, zeitgeber times ZT1 to 24). Most sleep events occurred at night (Fig. 7A-B and Fig. S4, zeitgeber times ZT12 to 24). This pattern was disrupted in *sesB^org^* and *sesB^9Ed-1^* animals, which experienced increased awake-sleep transitions throughout the whole day (Fig. 7A-B, zeitgeber times ZT1 to 24). *sesB* deficient animals slept more (Fig. 7C-D and G), a phenotype that was evident during the day (Fig. 7A-B, zeitgeber times ZT1 to 12 and Fig 7E) and night (Fig. 7A-B, zeitgeber times ZT12 to 24 and Fig 7F). The sleep increase phenotype observed in *sesB* alleles was selectively phenocopied only by glutamatergic neuron-specific sesB RNAi (Fig. 7H, Vglut driver). Neither glial-specific (Fig. 7I, repo driver) nor dopaminergic neuron-specific sesB RNAi elicited any sleep phenotypes (Fig. 7J, Ddc driver). These *sesB*-dependent phenotypes were in sharp contrast with the *sea*-dependent traits in two key aspects. First, sea RNAi decreased, rather than increased, total sleep duration, but only during the light period (Fig. S4A-B, zeitgeber times ZT1 to 12 and compare Fig. S4C-D with E-F). Second, this light-selective phenotype was only induced by downregulation of dSLC25A1-sea in dopaminergic neurons but not in glutamatergic neurons (Fig. S4 compare C-D with I-J). These results demonstrate that partial loss of function in SLC25A1 or SLC25A4 produce neuronal-cell type specific alterations of sleep. The synaptic and behavioral phenotypes support the notion that alterations in the expression of SLC25A1 and SLC25A4 participate either in pathogenesis and/or phenotypic expression of neurodevelopmental disorders.

**Fig. 7.**
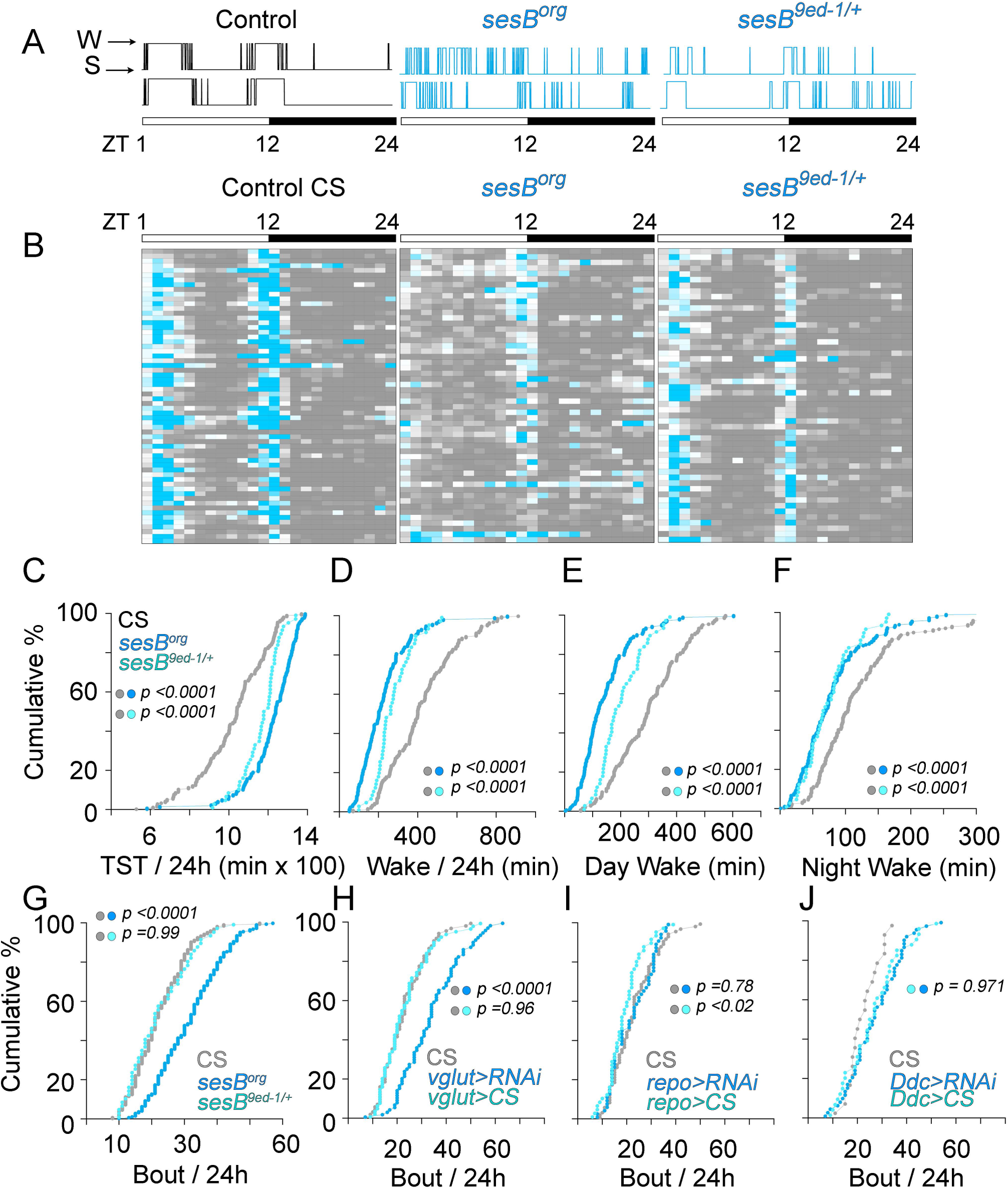
The *Drosophila* SLC25A4 Orthologue, *sesB,* is Required in Glutamatergic Neurons for Sleep. A) Individual hypnograms of two Canton S control and two *sesB* mutant flies illustrate sleep-wake activity patterns across the 12:12 hour light (zeitgeber times ZT1 to 12) and dark (zeitgeber times ZT12 to 24) periods. B) Heat map of sleep-wake activity (gray and teal, respectively) in Canton S control (n=229), *ses^org^* (n=234), and *sesB^9ed-1/+^* (n=53) depict the activity for each animal averaged across one hour bins. Each column is one zeitgeber hour and each row an animal. C-G) Probability plots of sleep parameters per 24 hours (C, D and G) or 12 hours light/darkness periods (E and F) from animals depicted in B. TST is total sleeping time. H) The number of sleep bouts per 24 hours is increased by *sesB* RNAi targeted to glutamatergic neurons (CS=78, VGlut>CS=72, VGlut>RNAi=82 animals) but neither in I) glial cells (CS=78, repo>CS=53, repo>RNAi=59 animals), nor J) catecholaminergic neurons (CS=21, Ddc>CS=37, Ddc>RNAi=56 animals). C-J) p values were estimated with the Kolmogorov–Smirnov test.

## Discussion

We identified mitochondrial pathways as statistically prioritized ontological terms in the 22q11.2 and the *Df(16)A+/-* proteomes. Our results recapitulate a previous proteomic study of *Df(16)A+/-* brains which is also enriched in mitochondrial targets (Fig. S5) (Wesseling et al., 2017). Here, we expanded this prior study by demonstrating first, that mitoproteomes are affected in neuronal and non-neuronal cells carrying this microdeletion. Second, we identified a novel interaction between the inner mitochondrial transporters SLC25A1 and SLC25A4; this physical, functional, and genetic interaction forming a high connectivity hub downstream to the 22q11.2 microdeletion (Fig. 4 and S3). The integrity of these mitochondrial transporters is required for normal synapse function and behavior as demonstrated by *Drosophila* hemideficiencies in dSLC25A4-*sesB* (Figs. 5-7). We further curated a SLC25A1-SLC25A4 interactome using comprehensive *in silico* tools and found that the expression of SLC25A transporters belonging to this interactome is altered in neurons from schizophrenia cases where the genetic risk factor is other than the 22q11.2 microdeletion (Fig. 4E-G). Our systems analysis of the 22q11.2 and the *Df(16)A+/-* proteomes intersects with studies where changes in mitochondrial ontologies strongly associate with psychiatric disease (Gandal et al., 2018) and with reports of alterations in mitochondrial transcripts, protein composition, function, and morphology in brains from patients with psychotic disorders (Enwright Iii et al., 2017; Middleton et al., 2002; Norkett et al., 2017; Rosenfeld et al., 2011). We propose that the mitochondrion, in particular components of the SLC25A1-SLC25A4 interactome, are *<u>a</u>* link in the chain of events connecting genetic risk factors with the expression of psychiatric phenotypes.

The robustness of our gene ontology conclusions is founded on the strong agreement between the 22q11.2 and the *Df(16)A+/-* proteomes. However, we deem necessary to discuss limitations inherent to the genealogical proteomic approach used here to generate the 22q11.2 proteome. First, the use of fibroblasts limits proteomic surveys to uncover systemic rather than neuronal-specific molecular phenotypes. Second, we observed that gene ontologies derived from different families are not overlapping despite the consistency of ontologies obtained from the same family yet with different quantitative proteomics methods (Fig. S1). Although we minimized the noise introduced by genetic variability among subjects by comparing proteomes within a family; noise introduced by variables like age, sex, cell passage, and epigenetic modifications due to possible drug use still contribute to our datasets. To circumvent these limitations, we reasoned that if 22q11.2 proteomes from different families contained a majority of 22q11.2-specific hits plus different sources of random noise, then the addition of these family-specific proteomes into one dataset should enrich 22q11.2-specific gene ontologies while degrading ontologies due to random noise. We empirically tested this idea by adding the 22q11.2 and the *Df(16)A+/-* proteomes, which resulted in improved statistical scores for the ontological categories associated to each one of these two datasets while maintaining their overall priority rank. In contrast, addition of a random gene dataset of increasing size progressively degraded statistical scores and/or ranking of microdeletion-specific ontological categories (Fig. S5C). These *in silico* data analyses support our approach of adding 22q11.2 family-specific proteomes to enrich gene ontologies affected by a genetic defect while diluting those generated by random noise. However, the best evidence supporting genealogical proteomics are the two independent *Df(16)A+/-* brain proteomes which validate of our studies. All three datasets, one in human and two in mouse, converge on similar ontological categories and rankings despite differences in tissues and species. The use of isogenic model systems is a way to circumvent random noise introduced by limited number of families in genealogical proteomics. Alternatively, either increasing the number of families analyzed or using different biological samples from the same family (fibroblasts, lymphoblasts, and IPSCs) should minimize the effects of noise on a dataset due to unforeseen variables or independent variables out of experimental reach.

The 22q11.2 locus encodes seven proteins contained in the Mitocarta 2.0 mitoproteome (COMT, MRPL40, PI4K, PRODH, SLC25A1, SNAP29, and TXNRD2) (Calvo et al., 2016; Pagliarini et al., 2008). The mitoproteome of 22q11.2 mutant cells and *Df(16)A+/-* brain likely reflect the collective effect of some or all these seven hemideficient genes (Devaraju et al., 2017; Devaraju and Zakharenko, 2017). We argue that these seven genes may not be the only 22q11.2 loci contributing to the alterations in the mitoproteome. For example, the DGCR8 gene, controlling microRNA-production, and seven miRNAs present in the 22q11.2 chromosomal segment could modulate the mitochondrial proteome acting both in nuclear and mitochondrial encoded RNAs (Bandiera et al., 2011; Chan et al., 2009; Minones-Moyano et al., 2011; Stark et al., 2008; Zhang et al., 2014). The seven 22q11.2 genes which are part of Mitocarta 2.0 are differentially expressed in different brain regions and cell types. Thus, their expression could influence the extent and quality of changes in mitochondrial proteomes from different cell types and brain regions in normal and disease states. We found that the stoichiometry of the mitoproteome or mitotranscriptome is different between two brain regions in normal mouse brain and between different cell types within *Drosophila* mushroom bodies (Fig. 2 and S2). We believe these regional and cellular differences in mitochondrial composition stoichiometry are consequential as the functional impact of downregulating dSLC25A1-dSLC25A4 on *Drosophila* sleep patterns depends on whether glutamatergic or catecholaminergic cells are targeted. It remains to be confirmed whether these effects are phenocopied by cell type-specific downregulation or knock-out of other components of the SLC25A1-SLC25A4 interactome in vertebrate and invertebrate brains. However, it is reasonable to propose that differences in mitochondrial composition stoichiometry in different neuronal cell types could explain why, of the SLC25A transcripts analyzed, SLC25A3 was the most affected in layer 3 parvalbumin cells and SLC25A4 the most affected in layers 3 and 5 pyramidal cells in subjects with schizophrenia (Fig. 4E).

22q11.2 microdeletion syndrome increases the risk of developing schizophrenia or Parkinson’s disease by 20-fold (Bassett and Chow, 2008; Bassett et al., 2000; Butcher et al., 2013; Butcher et al., 2017; Hodgkinson et al., 2001; Mok et al., 2016; Zaleski et al., 2009). This observation prompted us to ask about the identity of candidate pathways capable of contributing to both psychiatric and neurodegenerative phenotypes. Complex I and other respiratory complex subunits are prominently represented in the 22q11.2 and *Df(16)A+/-* proteomes and the SLC25A1-SLC25A4 interactome (Figs. 4 and S3). Respiratory chain complexes could contribute to the expression of psychiatric and/or neurodegenerative pathologies. Our contention is founded on the capacity of complex I chemical inhibitors to either cause Parkinson’s-like phenotypes (MPP+ and rotenone) or ameliorate psychosis symptoms (haloperidol, chlorpromazine, risperidone) (Burkhardt et al., 1993; Elmorsy and Smith, 2015; Modica-Napolitano et al., 2003; Prince et al., 1997; Rosenfeld et al., 2011). Moreover, primarily mitochondrial diseases that affect the activity of the respiratory chain complexes, such as Leigh syndrome, can cause neurodegeneration and psychiatric symptoms (Anglin et al., 2012a; Anglin et al., 2012b; DiMauro and Schon, 2008; Sheng and Cai, 2012). While still speculative, we put forward a testable model where tuning the activity of mitochondrial respiratory chain complexes by modifying the function of the diverse components within SLC25A1-SLC25A4 interactome, either by genetic and/or environmental factors, alters the expression of psychiatric and/or neurodegenerative phenotypes.

## Acknowledgements

This work was supported by grants from the National Institutes of Health R56 MH111459 to VF and Emory Catalyst Grant, Fondecyt-Chile grant 1171014 to GMR, R01HL108882 to SMC, R01MH097879 to JAG. We are indebted to the Faundez lab members for their comments. Stocks obtained from the Bloomington *Drosophila* Stock Center (NIH P40OD018537) were used in this study.

## STAR METHODS

### CONTACT FOR REAGENT AND RESOURCE SHARING

Further information and requests for resources and reagents should be directed to and will be fulfilled by the Lead Contact, Victor Faundez.

### EXPERIMENTAL MODEL AND SUBJECT DETAILS

#### Cell lines and Culture Conditions

Pedigrees of Ch22q11.2 fibroblasts were obtained from RUCDR Infinite Biologics repository (RUID: MH0162519, RUID: MH0162508, MH0162509, RUID: MH0162499, RUID: MH0162510, RUID: MH0162511, RUID: MH0162626, RUID: MH0162636, RUID: MH0162627, RUID: MH0162628, RUID: MH0162673, MH0162674, RUID: MH0162675, MH0162676, RUID: MH0162677, RUID: MH0162678). The fibroblasts were grown according to supplier recommendations in DMEM (Corning, 10-013- CV) media supplemented with 15% fetal bovine serum (FBS) (Atlanta Biologicals, S12450) and 100 μg/ml penicillin and streptomycin (Hyclone, SV30010) at 37°C in 5% CO_2_. SH-SY5Y cells (ATCC, CRL-2266; RRID: CVCL_0019) were cultured in DMEM media supplemented with 10% fetal bovine serum and 100 μg/mL penicillin and streptomycin at 37°C in 10% CO_2_. The SHSY-5Y cells were stably transfected either with a control empty vector (Genecopoeia, EX-NEG-Lv102) or ORF expression clone for N terminally tagged FLAG-SLC25A1 (Genecopoeia, EX-A1932-Lv1020GS) and grown in a selection media containing DMEM media supplemented with 10%FBS and Puromycin 2ug/ml (Invitrogen, A1113803). HEK293-Flp-In-pCDNA5/FRT-CNAP-Ant1/Ant2 (SLC25A4/SLC25A5) cells previously described (Lu et al., 2017). The cells were grown in DMEM media with 10%FBS and 100ug/ml Hygromycin (Invitrogen, 10687010). HAP1 cell lines – Control (C631), SLC25A1 knockout cell lines (HZGHC001753c003 and HZGHC001753c010) and SLC25A4 knockout cell line (HZGHC000778c011) were obtained from Horizon. HAP1 cells were cultured in IMDM media (Lonza, 12-722F) supplemented with 10% FBS and 100ug/ml penicillin and streptomycin at 37°C in 10% CO_2_.

#### Drosophila Husbandry and Stocks

Drosophila stocks were reared at 25°C in a humidified incubator (Shel Lab SR120PF) with a 12hr light/dark cycle and fed standard molasses food (900mL milli-Q water+48g active dry yeast+120g cornmeal+9g agar+120g molasses+2.4g tegosept+9mL propionic acid). The following stocks were used: w[1118] (#3605), C155-GAL4 (P{w[+mW.hs]=GawB}elav[C155] #458), Ddc-GAL4 (w[1118];P{w[+mC]=Ddc-GAL4.L}Lmpt[4.36] #7009) were obtained from the Bloomington Drosophila Stock Center. Gal4 lines used: c739 (α/βKC), NP1131 (γKC), R27G01 (MBON-γ5β’2a), R71D08(V2), G0431 (DAL). R27G01 (49233), G0239 (12639), G0431 (12837) and UAS-2xeGFP(6874) were ordered from the Bloomington Stock Center. NP1131-Gal4 was ordered from DGRC Stock center. R71D08 was a kind gift from Dr. H. Tanimoto. 3-86-Gal4 was a kind gift from Dr. U. Heberlein. c739-Gal4 was a kind gift from Dr. A. Sehgal. All Gal4 lines were crossed with UAS-2xeGFP to allow for cell harvesting.

#### Human Subjects

Seventy-seven patients with a molecularly confirmed diagnosis of 22q11DS and 50 unaffected, demographically matched healthy controls who were part of an ongoing longitudinal study at the University of California, Los Angeles were included in the current analyses. 22q11DS participants were recruited from posts to 22q11DS/Velocardiofacial online foundations and flyers through contacts with local craniofacial or genetics clinics. Controls were recruited from flyers posted at local schools and community centers. The study was approved by the UCLA Institutional Review Board and performed in accordance with the Declaration of Helsinki. All subjects or their legal guardians provided written informed consent and/or assent. This cohort has been previously published (Jalbrzikowski et al., 2015)

All data from the studies performed in postmortem human brain tissue have been previously published (Arion et al., 2015; Enwright Iii et al., 2017), and all methods and materials descriptions and data are publicly available (Arion et al., 2015; Enwright Iii et al., 2017).

### METHOD DETAILS

#### Antibodies

Antibodies used for immunoblots were as follows – SLC25A1 (Proteintech, 15235-1-AP; RRID: AB_2254794), SLC25A4 (1F3F11, a gift from the Claypool laboratory, Johns Hopkins), β-Actin (Sigma-Aldrich, A5441; RRID: AB_476744), HSP90 (BD Biosciences, 610418; RRID: AB_397798), TFRC (Invitrogen, 13-6800; RRID: AB_86623), FLAG (Bethyl, A190-102A; RRID: AB_67407). Blotting secondary antibodies were against mouse or rabbit conjugated to HRP (ThermoFisher Scientific, A10668; RRID: AB_2534058 and G21234; RRID: AB_2536530).

#### Cell Lysis and Immunoprecipitation

Cells intended for immunoprecipitation (Control HAP1 cells, HAP1 with SLC25A1/SLC25A4 knockdowns, SHSY-5Y empty vector or SHSY-5Y transfected withFLAG- SLC25A1 or HEK293-Flp-In-pCDNA5/FRT-CNAP-Ant1/Ant2 cells) were placed on ice, rinsed twice with ice cold PBS (Corning, 21-040-CV) containing 0.1mM CaCl2 and 1.0mM MgCl2. The cells were then rinsed twice with PBS and lysed in buffer A (150 mM NaCl, 10 mM HEPES, 1 mM EGTA, and 0.1 mM MgCl_2_, pH 7.4) with 0.5% TritonX-100 and Complete anti-protease (Roche, 11245200). Cells were scraped from the dish, placed in Eppendorf tubes, followed by incubation for 30 minutes on ice. Cell homogenates were then centrifuged at 16,100×*g* for 10 minutes and the clarified supernatant was recovered. Protein concentration determined using the Bradford Assay (BioRad, 5000006). For immunoprecipitation, 500 μg of protein extract was incubated with 30 microliters Dynal magnetic beads (Invitrogen, 110.31) coated with antibody, and incubated for 2 hours at 4°C. In some cases, immunoprecipitations were done in the presence of the antigenic 3X FLAG peptide (340μM) (Sigma, F4799) as a control. The beads were then washed 4-6 times with buffer A with 0.1% TritonX-100. Proteins were eluted from the beads with Laemmli buffer. Samples were resolved by SDS-PAGE and contents analyzed by immunoblot described below.

#### Quantitative Mass Spectrometry

##### Stable Isotope Labeling of Amino Acids (SILAC)

Ch22q11.2 fibroblasts were labeled using published protocols. Cells were cultured in DMEM with either “light” unlabeled arginine and lysine amino acids (R0K0) (Dundee Cell Products, LM014) “medium” 13C- and 15N-labeled arginine, and 13C- and 15N-labeled lysine amino acids (R6K4) (Dundee Cell Products, LM016) or “heavy” 13C- and 15N-labeled arginine, and 13C- and 15N-labeled lysine amino acids (R10K8) (Dundee Cell Products, LM015) supplemented with 15% FBS (Dundee Cell Products, DS1003) and 100 μg/ml penicillin and streptomycin. Each cell line was grown for seven passages allowing maximum incorporation (at least 97.5%) of the amino acids in the total cellular pool. Cellular lysate samples were prepared, as described in the previous section. Quantitative mass spectrometry was performed as described previously using the services of MS Bioworks and the Emory Integrated Proteomics Core.

The SILAC labeled samples were pooled 1:1:1 and 20μg of this mix was resolved on a 4-12% Bis-Tris Novex mini-gel (Invitrogen) using the MOPS buffer system. The gel was stained with Coomassie and the lanes excised into 40 equal sections using a grid. Gel pieces were robotically processed (ProGest, DigiLab) by first washing with 25mM ammonium bicarbonate (ABC) followed by acetonitrile, followed by reduction with 10mM dithiothreitol at 60°C, alkylation with 50mM iodoacetamide at room temperature. Pieces were digested with trypsin (Promega) at 37°C for 4h and quenched with formic acid. The supernatant was analyzed directly without further processing. Gel digests were analyzed by nano LC/MS/MS with a Waters NanoAcquity HPLC system interfaced to a ThermoFisher Q Exactive. Peptides were loaded on a trapping column and eluted over a 75μm analytical column at 350nL/min; both columns were packed with Jupiter Proteo resin (Phenomenex). The mass spectrometer was operated in data-dependent mode, with MS and MS/MS performed in the Orbitrap at 70,000 FWHM resolution and 17,500 FWHM resolution, respectively. The fifteen most abundant ions were selected for MS/MS. Data were processed through the MaxQuant software 1.4.1.2 (www.maxquant.org) which served the following functions: 1. Recalibration of MS data. 2. Filtering of database search results at the 1% protein and peptide false discovery rate (FDR). 3. Calculation of SILAC heavy:light ratios. Data were searched using a local copy of Andromeda with the following parameters: Enzyme: Trypsin. Database: Swissprot (concatenated forward and reverse plus common contaminants). Fixed modification: Carbamidomethyl (C). Variable modifications: Oxidation (M), Acetyl (Protein N-term), ^13^C_6_/^15^N_2_ (K), ^13^C6/^15^N_4_ (R), ^4^H_2_ (K), ^13^C_6_ (R). Fragment Mass Tolerance: 20ppm. Pertinent MaxQuant settings were: Peptide FDR 0.01. Protein FDR 0.01. Min. peptide Length 7. Min. unique peptides 0. Min. ratio count 2. Re-quantify TRUE. Second Peptide TRUE.

###### Label-free Quantitation (LFQ) cellular preparation

Cells were grown in 10cm dishes to 85-90% confluency. On the day of the experiment the cells were placed on ice and washed 3 times with PBS supplemented with 10mM EDTA (Sigma, 150-38-9) for 3 minutes each. After the third wash, the cells were incubated with PBS and 10mM EDTA for 30 minutes on ice. Cells were then lifted with mechanical agitation using a 10ml pipette and collected in a 15ml falcon tube. Cells were then spun at 800xg for 5 minutes at 4°C. The supernatant was then aspirated out and the remaining pellet was washed with ice cold PBS. The resuspended cells were then centrifuged at 16,100×*g* for 5 min. The supernatant was discarded and the resulting pellet was immediately frozen on dry ice for at least 5 minutes and stored at - 80°C for future use.

Cell pellets were lysed in 200ul of urea lysis buffer (8M urea, 100 mM NaH2PO4, pH 8.5), supplemented with 2 uL (100x stock) HALT protease and phosphatase inhibitor cocktail (Pierce). Lysates were then subjected to 3 rounds of probe sonication. Each round consisted of 5 seconds of activation at 30% amplitude and 15 of seconds of rest on ice. Protein concentration was determined by bicinchoninic acid (BCA) analysis and 100 ug of each lysate was aliquoted and volumes were equilibrated with additional lysis buffer. Aliquots were diluted with 50mM ABC and was treated with 1mM DTT and 5mM IAA in sequential steps. Both steps were performed in room temperature with end to end rotation for 30 minutes. The alkylation step with IAA was performed in the dark. Lysyl endopeptidase (Wako) was added at a 1:50 (w/w) enzyme to protein ratio and the samples were digested for overnight. The following morning, a 50ug aliquot was taken out, acidified to a final concentration of 1% formic acid and stored. Trypsin (Promega) was added to the residual 50ug aliquot at a 1:100 (w/w) and digestion was allowed to proceed overnight again. Resulting peptides from both digestions rounds were desalted with a Sep-Pak C18 column (Waters).

Dried peptide fractions were resuspended in 100 μl of peptide loading buffer (0.1% formic acid, 0.03% trifluoroacetic acid, 1% acetonitrile). Peptide mixtures were separated on a self-packed C18 (1.9 μm Dr. Maisch, Germany) fused silica column (25 cm × 75 μm internal diameter; New Objective) by mass spectrometer platforms: 1) Dionex Ultimate 3000 RSLCNano coupled to a Fusion orbitrap tribrid mass spectrometer (ThermoFisher Scientific) and 2) Water’s NanoAcquity coupled to a Q-Exactive Plus hybrid mass spectrometer (ThermoFisher Scientific). For the Fusion system, 2ul was loaded and elution was carried out over a 140 min gradient at a rate of 300 nl/min with buffer B ranging from 3% to 99% (buffer A: 0.1% formic acid in water, buffer B: 0.1% formic in acetonitrile). The mass spectrometer cycle was programmed to collect at the top speed for 5 s cycles consisting of 1 MS scan (400-1600 m/z range, 200,000 AGC, 50 ms maximum ion time) were collected at a resolution of 120,000 at m/z 200 in profile mode followed by ion trap collected HCD MS/MS spectra (0.7 m/z isolation width, 30% collision energy, 10,000 AGC target, 35 ms maximum ion time). Dynamic exclusion was set to exclude previous sequenced precursor ions for 20 s within a 10 ppm window. Precursor ions with +1 and +8 or higher charge states were excluded from sequencing. For the Q-Exactive Plus system, 4 ul was loaded and elution was carried out over a 140 min gradient at a rate of 250nl/min with buffer B ranging from 3 to 80% ACN. The mass spectrometer was set to acquire 1 MS scan (70,000 resolution at 200 m/z in profile mode, 300-1800 m/z range, 1,000,000 AGC, 100 ms maximum ion time) followed by at most 10 MS/MS scans(17,500 resolution at 200 m/z, 2.0 m/z isolation width with an offset of 0.5 m/z, 50,000 AGC, 50 ms maximum ion time). Dynamic exclusion was for 30 s with a 10 ppm window.

All spectra from both platforms were loaded into Maxquant (version 1.5.2.8) and searched against a database downloaded from the NCBI’s REFSEQ (version 54) with common contaminants appended. Search parameter included fully tryptic (or lysyl endopeptidase) cleavage, variable modifications for Protein N-terminal acetylation and methionine oxidation, static modifications for cysteine carbamindomethyl, 20 ppm precursor mass tolerance, 0.5 Da for ion trap and .05 Da for Orbitrap product ion tolerances, FDR at 1% for all levels including protein, peptide and psm.

##### Tandem Mass Tagging (TMT)

Cell pellets were lysed, reduced, alkylated and digested similarly as with the LFQ protocol with the only differences being that 50mM TEAB was used for dilution and only Lysyl endopeptidase was used for digestion. An aliquot equivalent to 10 ug of total protein was taken out of each sample and combined to obtain a global internal standard (GIS) use later for TMT labeling.

TMT labeling was performed according to the manufacturer’s protocol. Briefly (Ping et al., 2018), the reagents were allowed to equilibrate to room temperature. Dried peptide samples (90 μg each) were resuspended in 100 μl of 100 mm TEAB buffer (supplied with the kit). Anhydrous acetonitrile (41 μl) was added to each labeling reagent tube and the peptide solutions were transferred into their respective channel tubes. The reaction was incubated for 1 h and quenched for 15 min afterward with 8 μl of 5% hydroxylamine. All samples were combined and dried down. Peptides were resuspended in 100 μl of 90% acetonitrile and 0.01% acetic acid. The entire sample was loaded onto an offline electrostatic repulsion–hydrophilic interaction chromatography fractionation HPLC system and 40 fractions were collected over a time of 40 min. The fractions were combined into 10 and dried down. Dried peptide fractions were resuspended in 100 μl of peptide loading buffer (0.1% formic acid, 0.03% trifluoroacetic acid, 1% acetonitrile). Peptide mixtures (2 μl) were separated on a self-packed C18 (1.9 μm Dr. Maisch, Germany) fused silica column (25 cm × 75 μm internal diameter; New Objective) by a Dionex Ultimate 3000 RSLCNano and monitored on a Fusion mass spectrometer (ThermoFisher Scientific). Elution was performed over a 140 min gradient at a rate of 300 nl/min with buffer B ranging from 3% to 80% (buffer A: 0.1% formic acid in water, buffer B: 0.1% formic in acetonitrile). The mass spectrometer cycle was programmed to collect at the top speed for 3 s cycles in synchronous precursor selection mode (SPS-MS3). The MS scans (380-1500 m/z range, 200,000 AGC, 50 ms maximum ion time) were collected at a resolution of 120,000 at m/z 200 in profile mode. CID MS/MS spectra (1.5 m/z isolation width, 35% collision energy, 10,000 AGC target, 50 ms maximum ion time) were detected in the ion trap. HCD MS/MS/MS spectra (2 m/z isolation width, 65% collision energy, 100,000 AGC target, 120 ms maximum ion time) of the top 10 MS/MS product ions were collected in the Orbitrap at a resolution of 60000. Dynamic exclusion was set to exclude previous sequenced precursor ions for 30 s within a 10 ppm window. Precursor ions with +1 and +8 or higher charge states were excluded from sequencing.

MS/MS spectra were searched against human database from REFSEQ (version 54) and Uniprot (downloaded on 03/06/2015) with Proteome Discoverer 1.4 and 2.0 (ThermoFisher Scientific), respectively. Methionine oxidation (+15.9949 Da), asparagine, and glutamine deamidation (+0.9840 Da) and protein N-terminal acetylation (+42.0106 Da) were variable modifications (up to 3 allowed per peptide); static modifications included cysteine carbamidomethyl (+57.0215 Da), peptide n terminus TMT (+229.16293 Da), and lysine TMT (+229.16293 Da). Only fully cleaved Lysyl endopeptidase peptides were considered with up to two miscleavages in the database search. A precursor mass tolerance of ±20 ppm and a fragment mass tolerance of 0.6 Da were applied. Spectra matches were filtered by Percolator to a peptide-spectrum matches false discovery rate of <1%. Only razor and unique peptides were used for abundance calculations. Ratio of sample over the GIS of normalized channel abundances were used for comparison across all samples.

#### Electrophoresis and Immunoblotting

For western blot, lysate was reduced and denatured in Laemmli buffer containing SDS and 2-mercaptoethanol and heated for 5 minutes at 75°C. Equal amounts of cellular lysates were loaded onto 4-20% Criterion gels (BioRad, 5671094) for electrophoresis and transferred to PVDF (Millipore, IPFL00010) using the semi-dry transfer method. The PVDF membranes were blocked with Tris buffered saline containing 5% non-fat milk and 0.05% Triton X-100 (TBST), rinsed and incubated overnight in presence of appropriately diluted primary antibody in antibody base solution (PBS with 3% Bovine Serum Albumin, 0.2% Sodium Azide). Membranes were then washed multiple times in TBST and incubated in HRP conjugated secondary antibody diluted 1:5000 in the blocking solution above. Following multiple washes, the membranes were then exposed to Amersham Hyperfilm ECL (GE Healthcare, 28906839) with Western Lightning Plus ECL reagent (Perkin Elmer, NEL105001EA).

#### Cell Line RNA Extraction and Quantitative RT-PCR

RNA extraction for cells and tissues was done using Trizol Reagent (Invitrogen, 15596026) following the published protocol. Total amount, concentration and purity of RNA were determined using the BioRad SmartSpec Plus Spectrophotometer. First strand synthesis was completed using the Superscript III First Strand Synthesis System Kit (Invitrogen, 18080-051) utilizing 5 μg total RNA per reaction and random hexamer primers following manufactures protocol. RT-PCR was done with 1μl cDNA from first strand synthesis in LightCycler 480 SYBR Green I Master (Roche, 04707516001) according to manufactures protocol on a LightCycler 480 Instrument with 96-well format. RT-PCR protocol included an initial denaturation at 95° C for 5min, followed by 45 cycles of amplification with a 5 second hold at 95°C ramped at 4.4°C/s to 55°C. Temperature was then held for 10 seconds at 55°C and ramped up to 72°C at 2.2°C/s. Temperature was held at 72°C for 20 seconds were a single acquisition point was collected and then ramped at 4.4°C/s to begin the cycle anew. A melting curve was collected following amplification. The temperature was then held at 65° for 1 min and ramped to 97°C at a rate of 0.11°C/s. Five acquisition points were collected per °C. Primers were designed using the IDT Real Time qPCR Assay Entry site using site recommended parameters. Primers were obtained from Sigma Custom DNA Oligo service. Melting curves were used to confirm primer specificity to single transcripts. A primer list is provided in Table X. For quantification, standard curves for each primer were applied to all samples using LightCycler 480 software. Ratios of experimental to control samples, normalized to reference genes, are reported.

#### Drosophila Neuromuscular Microcopy

Neuromuscular junction staining was performed using late third instar larvae. Larval body wall dissections using a dorsal incision were performed with 10mM cell culture dishes partially filled with charcoal infused sylguard, microdissection pins, forceps and microdissection scissors. Drosophila were dissected using in standard Ca2+ free HL3 Ringer’s Solution (70mM NaCl, 5mM KCl, 21.5mM MgCl2, 10mM NaHCO3, 5mM Trehalose, 115mM Sucrose, 5mM BES with pH of 7.2-7.3), fixed using 4% paraformaldehyde for 45minutes-1hour at room temp, rinsed 10minutes with PBS-T (PBS+.15% Triton), incubated in FITC-HRP conjugate (MP Biomedicals #0855977) overnight at 4C. Rinses followed the next day in PBS-T at 3×1minute rinse then 3×10minute rinse and finished with a 3×1minute rinse in PBS. Larval body wall preparations were then placed on slides with a drop of Vectashield and coverslip. Nail polish was used to seal the edges of the coverslip in place and samples were stored at 4C until imaged. Confocal images were obtained using a Zeiss LSM 510 microscope and Zen 2009 software. NMJs from 6/7 muscles of the 3rd or 4th segments were identified and z-stack images collected with a continuous wave 458,488nm argon laser at 200mW. Z-stacks were converted to jpegs using FIJI software and blinded for bouton quantification.

#### Drosophila Electrophysiology

NMJ dissections of third instar, female larvae were performed in ice-cold, calcium-free HL-3 Ringer’s solution (70mM NaCl, 5mM KCl, 21.5mM MgCl_2_, 10mM NHCO_3_, 5mM Trehalose, 115mM Sucrose, 5 mM BES in ddH_2_0 with a pH of 7.2-7.3). After the dissection, the filleted preparation was rinsed twice in low-Ca^2+^ Ringer’s solution (70mM NaCl, 5mM KCl, 1mM CaCl_2_, 20mM MgCl_2_, 10mM NHCO_3_, 5mM Trehalose, 115mM Sucrose, 5 mM BES in ddH_2_0 with a pH of 7.2-7.3) and the low-Ca^2+^ Ringer’s solution was used throughout the remainder of the experiment. Motor axons were severed close to the ventral ganglion and were taken up into a borosilicate glass capillary suction electrode with a firepolished tip (Microforge MF-830, Narishige). Recording electrodes were prepared using borosilicate glass capillary tubes (1mm outer diameter and 0.58 internal diameter; A-M Systems) which were pulled to a fine tip (PN-3, Narishige) with 25-50 MOhm resistance and backfilled with 3M KCl. All recordings were obtained from muscle 6 in the second or third abdominal sections (A2 or A3, respectively). Stimulations were delivered using a Model 2100 Isoplated Pulse Stimulator (A-M Systems) and recordings were acquired with an Axoclamp 900A amplifier (Molecular Devices). pClamp 10 software (Molecular Devices; RRID:SCR_011323) was used to collect data and analyzed EJP amplitude and membrane potential and Mini Analysis Program (Synaptasoft; RRID:SCR_002184) was used to analyze mEJPS frequency and amplitude.

#### Drosophila Behavior (Sleep)

Female flies were collected under CO_2_ anesthesia within 72 hours of eclosure. Twenty-four hours later, flies were briefly cooled on ice to allow mouth pipetting of individual flies into polycarbonate tubes (5mm external diameter × 65mm; TriKinetics.com). One end of the tube contained a 5% (w/v) sucrose and 2% (w/v) agarose medium while the other end was sealed with parafilm perforated with an 18-gauge needle to allow air circulation. Tubes were placed in the *Drosophila* Acitivity Monitoring System (DAM2; TriKinetics) which was housed in a light-controlled cabinet with a 12h:12h light:dark cycle at room temperature.

Data was collected in 15 second intervals using the DAMSystem308 acquisition software (TriKinetics) and analysis was based upon 1-minute bins across 6 days of data collection, starting at Lights ON the day after the animals were placed in the tubes. Periods of inactivity lasting longer than 5 minutes were scored as sleep (Hendricks et al. 2000, Shaw et al. 2000) and sleep duration, bout number, and bout length were calculated using a custom created analysis in Excel. All genotypes were compared to Canton S. Each UAS- and Gal4- line was crossed to Canton S to verify that neither the presence of the transgenes nor the genetic background of these individual lines altered the sleep/wake phenotype. Sleep/wake phenotypes of the sesB mutants were assessed based upon homozygous populations of the hypomorphic sesB^org^ mutation and heterozygous populations of the lethal sesB^9Ed-1^mutation (sesB^9Ed-1^/FM7a;;).

#### Drosophila RNAseq library generation

##### Cell harvesting

GFP labeled cells were handpicked in vivo through suction into a pipette. Cells designated for sequencing were harvested into .5ul nuclease free water in the pipette tip and then the tip was broken into a 96 well PCR tube containing RNAse inhibitors and buffer as described by Clontech’s ultra low HV SMARTer Ultra Low RNAseq kit (Catalog# 634823) resulting in the lysing of cells without mechanical means. Amplification was performed following the Clontech Ultra-Low volume SMARter RNAseq Protocol. For the DAL neuron, the MBONα3 neurons, the MBON-γ5β’2a and MBON-β2β’2a neurons, 4 cells were pooled into each tube thus these samples contained cells from more than one fly. For the V2, α/βKCs and γKCs all cells were taken from one animal per sample. V2 samples contained 14 cells and the α/βKC and γKC samples contained about 100 cells. 15 rounds of PCR amplification were performed using the Clontech SMARTer Ultra low RNAseq Kit. For this work only cells collected from animals that had undergone unpaired odor and shock presentation were used.

Following amplification samples were selected if there was a peak around 7 kb and .4-2ng/ul of product between the range 400bp-10kb. Samples were then sheared using a Covaris LE220 sonicator to 200bp. The libraries were made using the IntegenX automated library prep system. The PrepX Illumina DNA library prep kit/ PrepX CHIPseq kit (WaferGen Biosystems Inc) was used with an amplification of 17-22 cycles. They were multiplexed using Bioo Scientific barcodes. Then cleaned using the IntegenX PCR cleanup kit. Libraries were run on the Illumina HiSeq2500, 12 samples per lane and each sample run across two lanes. Resulting in a sequencing depth of 30 million reads. Sequencing was all done single end.

##### Analysis of Sequencing Reads

FastQC (Andrews, 2012) was performed to remove samples of poor quality. Samples all contained a bias for polyA and T sequences. This was uniform across all samples and was removed from sequences prior to mapping. GC content was not flagged on samples used in the study. All mapping was performed using Princeton University’s Galaxy server running TopHat 2 with Bowtie2 (Kim et al.2013, (Langmead and Salzberg, 2012). The Ensembl build of the reference sequence (BDGF 5.25) and the GTF files were used and can acquired from iGenome (illumina). The aligned SAM/BAM file were processed using HTseq-count (Intersection mode - strict) (Anders et al., 2015). HTseq Counts output files and raw illumine read files are publicly available (GEO with accession number GSE4989). The HTseq counts compiled file is GSE74989_HTseqCountscompiledData.txt.gz

##### Calculating Normalized Gene Counts

The GSE74989_HTseqCountscompiledData.txt.gz data set was used for analysis. In R, all genes with counts less than 2 counts per million (8 counts) across all samples independent of cell-type were considered noise and removed from analysis. Gene counts were normalized using DESeq2 (Love et al., 2014) followed by a regularized log transformation. Genes with less than 2 counts per million within cell-type were recoded as zero. Principal component analysis was performed on this processed data set in R. R function prcomp was used to generate the principle components and gene loading values.

#### Drosophila Transcriptome Encoding Mitochondrial Proteins

All data was acquired from the GEO data set GSE74989 which is publicly available. From this data set only control animals were used to generate the figure and cell-type results. Thus 5 DAL samples, 5 V2 samples, 5 a/b KC samples, 5 gKC samples, 5 MBON b2b’2a, 4 MBON g5b’2a.

#### Human Postmortem RNA Analysis

All data from the studies performed in postmortem human brain tissue have been previously published. All tissue sample collection and RNA sequencing (RNASeq) details are publicly available (https://www.synapse.org/#!Synapse:syn2759792/wiki/). All tissue sample collection and microarray analysis details are described in detail (Arion et al., 2015; Enwright Iii et al., 2017), and the data are publicly available upon request.

#### 22q11DS Patient and Control RNAseq

RNA was extracted from whole blood using the PAXgene extraction kit (Qiagen), then stored at −80C for subsequent analysis. RNA quantity was assessed with Nanodrop (Nanodrop Technologies) and quality with the Agilent Bioanalyzer (Agilent Technologies). Gene expression profiling was performed using Illumina HT-12 v4 microarrays. 200 ng of total RNA were amplified, biotinylated and hybridized to Illumina Human V4-HT-12 Beadchips, including approximately 47,000 probes, following the manufacturer’s recommendations. Slides were scanned using Illumina BeadStation, and the signal was extracted by using Illumina BeadStudio software.

Raw data were analyzed using Bioconductor packages in the R statistical environment. Only samples with an RNA integrity number (RIN) of 7 or greater were included in the analyses. Gene expression variance was normalized using variance stabilized transformation. Quality assessment was performed by examining the inter-array biweight midcorrelation; samples more than 3 standard deviations from the mean were excluded. Batch effects were removed using ComBat. Differential gene expression analysis used the *limma* package in R to implement general linear model fit, with batch correction, age, sex, and RIN as covariates.

## QUANTIFICATION AND STATISTICAL ANALYSIS

Experimental conditions were compared using Synergy Kaleida-Graph, version 4.1.3 (Reading, PA) or Aabel NG2 v5 x64 by Gigawiz as specified in each figure.

## DATA AND SOFTWARE AVAILABILITY

“*The XXX have been deposited in the XXX under ID codes XXX and YYY.*”

---

**Supplementary Fig. 1.**
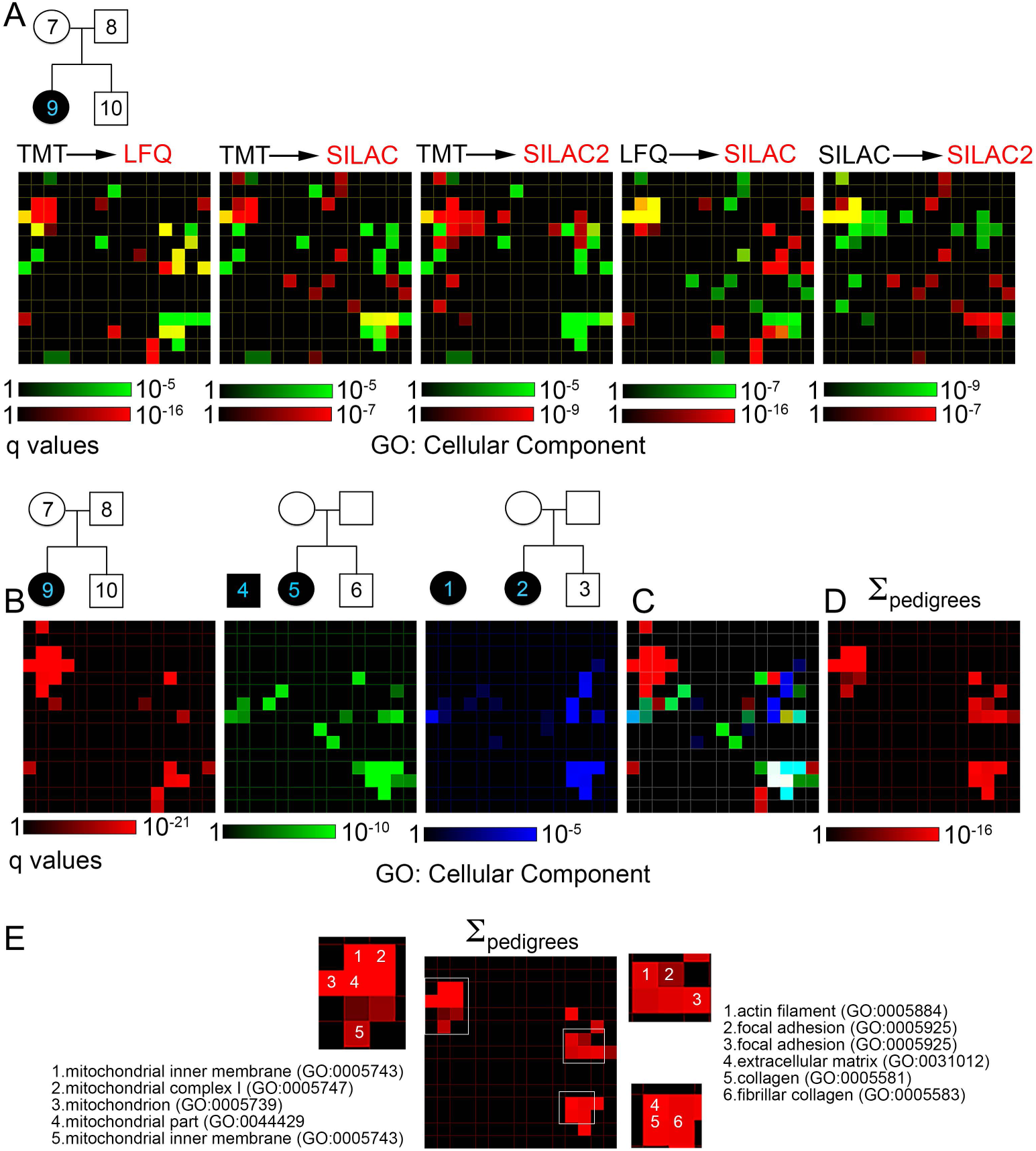
22q11.2 Microdeletion Genealogical Proteome Comparisons Among Pedigrees and Mass Spectrometry Quantitation Strategies. A-D) Cellular Component Gene ontologies (GO:CC) obtained using the ENRICHR engine. Data are depicted as canvases where every tile is occupied by an individual GO category whose p value significance is depicted by color intensity. A) First canvas to the left depicts a comparison for genealogical proteomes obtained in one pedigree by TMT (green) and to Label Free Quantification (LFQ, red). Other canvases in A show experiments comparing TMT with two independent SILAC experiments and combinations of TMT, LFQ and SILAC. Overlap of GO:CC terms is presented as yellow. B) Comparison of GO:CC terms obtained by genealogical proteomics from three pedigrees. C) Represents GO:CC term tiles overlapping among pedigrees in B). D) Shows gene ontology terms obtained by pooling into one dataset the proteomes from all pedigrees in B). E. Presents relevant GO terms in D. Individual pedigree and collective bioinformatics data can be found in Table S4.

**Supplementary Fig. 2.**
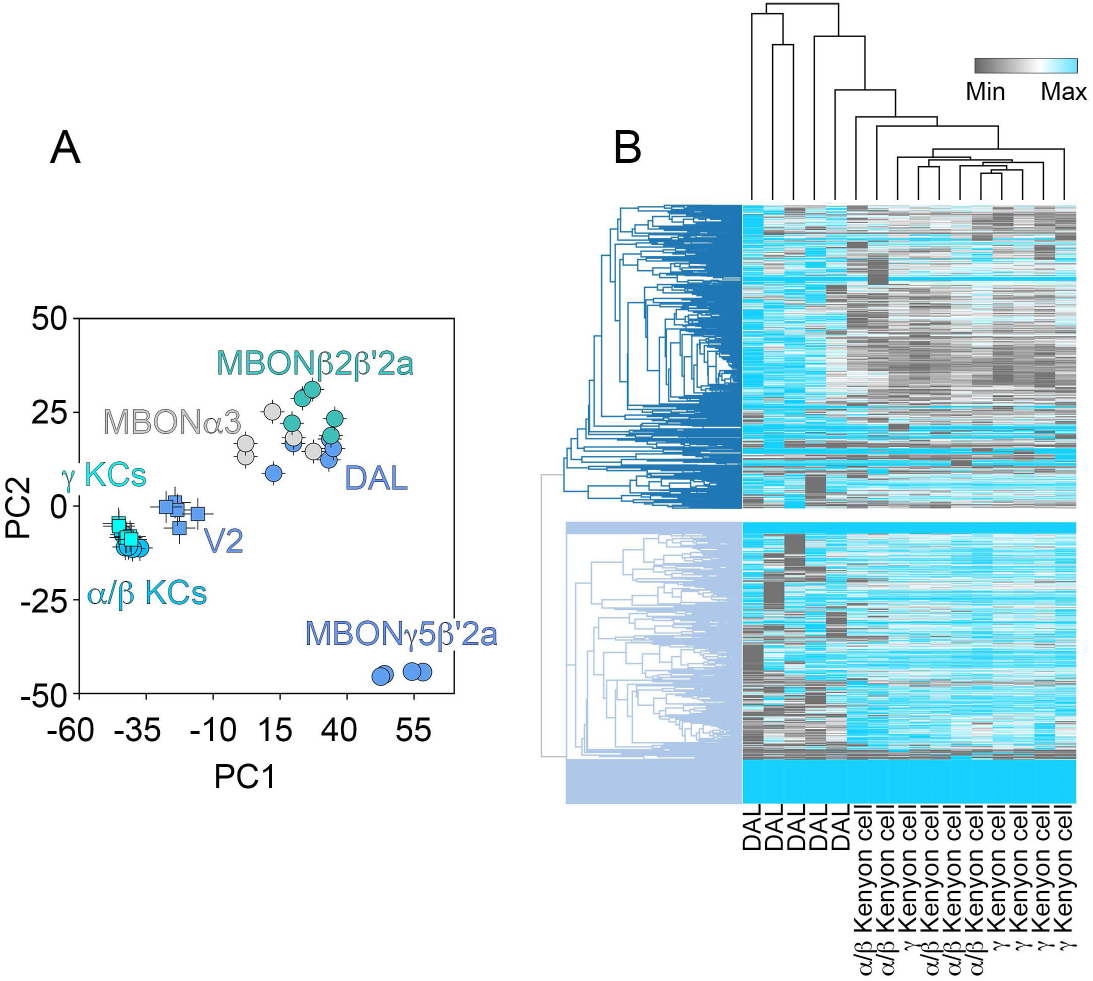
The Drosophila Transcriptome Encoding Mitochondrial Proteins is Cell Type Specific. A-B) mRNA from single neuron types isolated from *Drosophila* mushroom bodies were analyzed by RNAseq. The transcriptome encoding mitochondrial proteins, as defined by Chen et al (Chen et al., 2015), was analyzed by principal component analysis (A) and hierarchical clustering using 1-Pearson correlation clustering (B) of columns (cells) and rows (transcripts). Cell types were identified as in Crocker et al (Crocker et al., 2016). Note the robust segregation of Kenyon cells from other cell types by the expression of the transcriptome encoding mitochondrial proteins.

**Supplementary Fig. 3.**
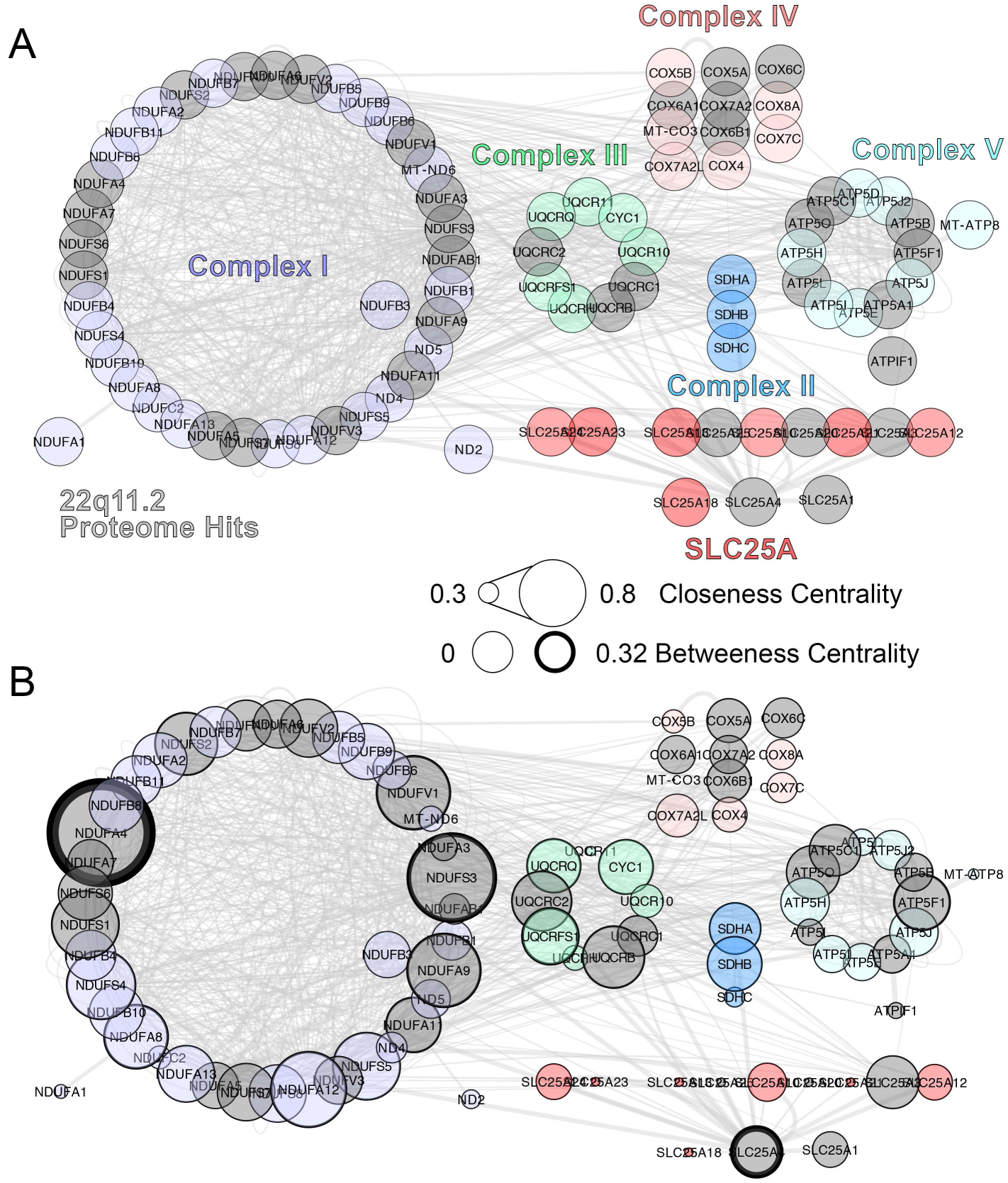
Comprehensive *in silico* Interactome of the SLC25A1 and SLC25A4 Transporters. A) Comprehensive *in silico* interactome of the SLC25A1 and SLC25A4 mitochondrial transporters. Complexes I to V of the respiratory chain as well as SLC25A transporter family members are color coded. All nodes colored gray represent hits in the 22q11.2 proteome. B) The comprehensive interactome was analyzed with graph theory to determine high connectivity nodes predictive of essential genes using the closeness and betweeness centrality coefficients. Note the high connectivity of SLC25A4 in the comprehensive interactome

**Supplementary Fig. 4.**
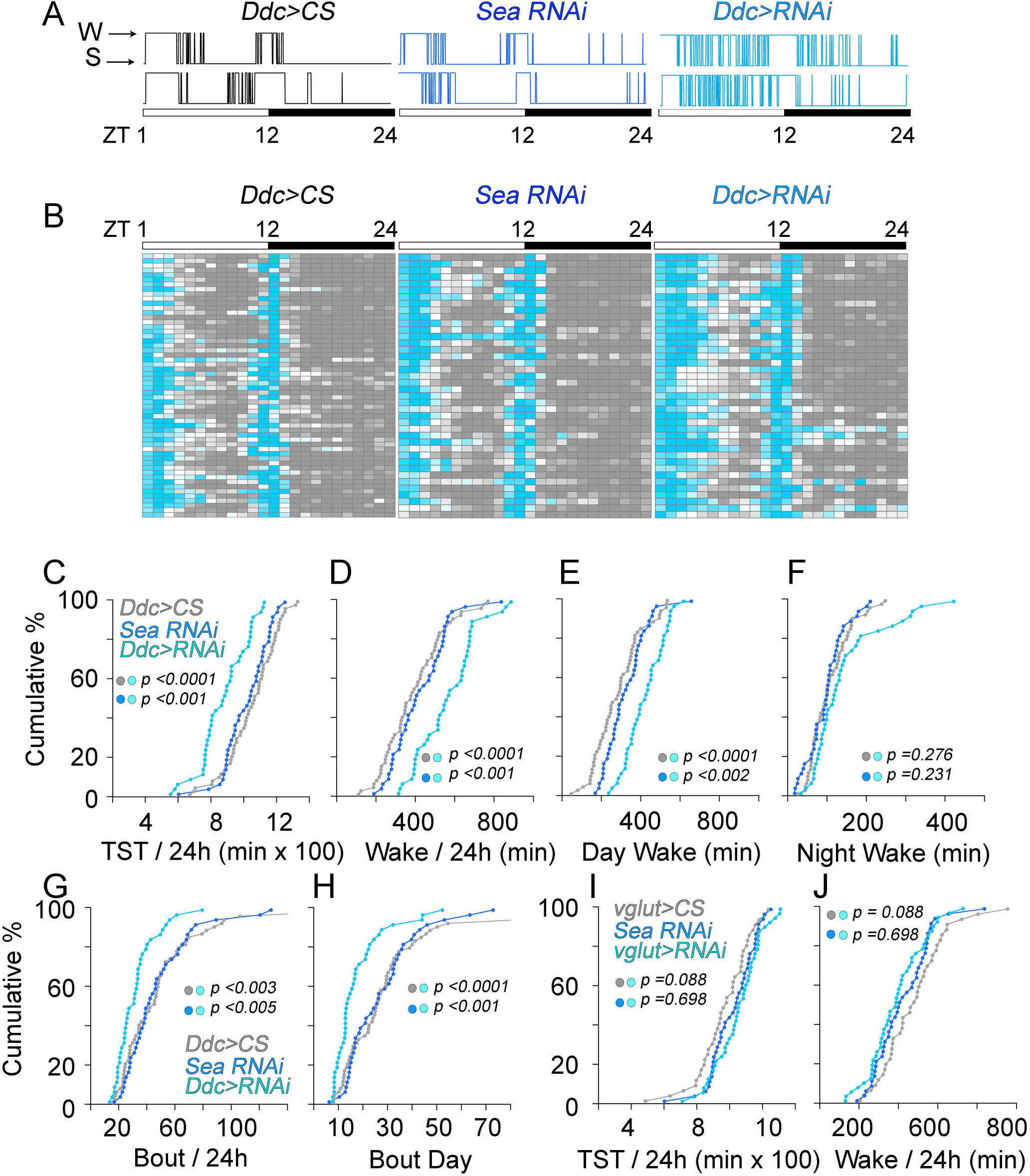
*Drosophila* SLC25A5 Orthologue *Sea* is Required in Catecholaminergic Neurons for Sleep. A) Individual hypnograms of Canton S control, *sea* RNAi controls, and catecholaminergic-specific *sea* RNAi (Ddc>RNAi) flies (n=2 each) illustrates sleep-wake activity patterns across the 12:12 hour light (zeitgeber times ZT1 to 12) and dark (zeitgeber times ZT12 to 24) periods. B) Heat maps of sleep-wake activity (gray and teal, respectively) in *Ddc driver* control (Ddc>CS, n=56), *sea RNAi* control (n=40), and catecholaminergic-specific *sea* RNAi animals (Ddc>RNAi, n=40) depict activity for each animal averaged across one hour bins. Each column is one zeitgeber hour and each row is one animal. C-H) Probability plots of sleep parameters per 24 hours (C, D and G) or 12 hours light/dark periods (E, F and H) from animals depicted in B. TST is total sleeping time. G-H) The number of sleep bouts is decreased in catecholaminergic-specific *sea* RNAi animals. No effect of glutamatergic-specific *sea* RNAi (VGlut>CS=38, sea RNAi=40, VGlut>RNAi= 44 animals) C-J) p values were estimated with the Kolmogorov–Smirnov test

**Supplementary Fig. 5.**
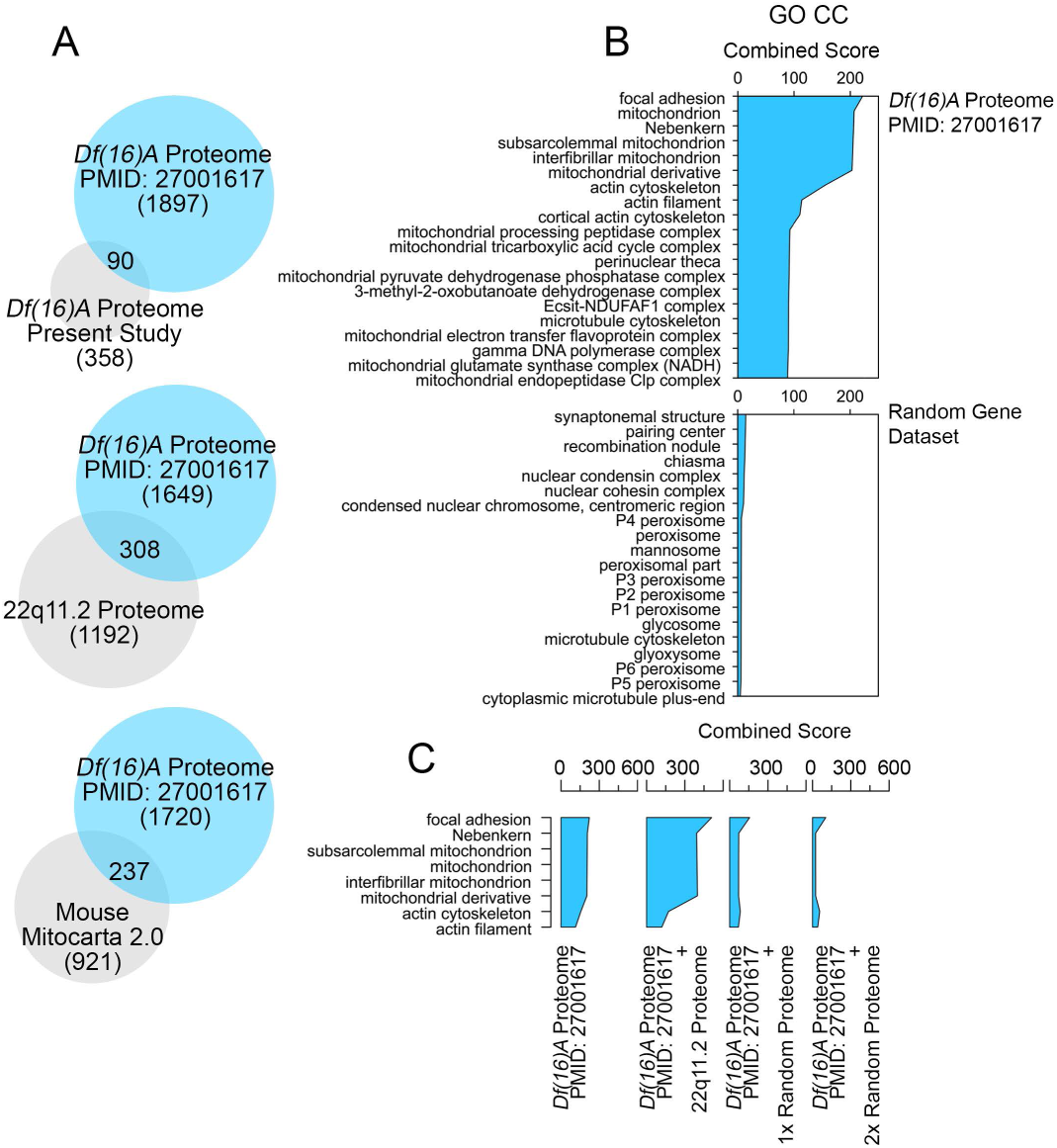
Comparative Bioinformatics of the 22q11.2 Proteome and Two Independent *Df(16)A*-/+ Brain Proteomes. A) Venn diagrams depict from top to bottom: a comparison of common hits between our *Df(16)A*-/+ brain proteome and the *Df(16)A*-/+ brain proteome reported by Wesseling et al. PMID: 27001617. The Wesseling *Df(16)A*-/+ brain proteome and our 22q11.2 proteome. The Wesseling *Df(16)A*-/+ brain proteome and the mouse Mitocarta 2.0 dataset. B) Cellular Component gene ontology analysis of GO CC generated with the ENRICHR engine using the Wesseling *Df(16)A*-/+ brain proteome dataset and a similarly sized random mouse gene dataset. Random gene list was generated with the engine [RandomGeneSetGenerator](http://www.molbiotools.com/randomgenesetgenerator.html). C) Cellular Component gene ontology analysis (GO CC) was performed with the ENRICHR engine using the Wesseling *Df(16)A*-/+ brain proteome dataset either by itself, or in combination with our 22q11.2 proteome, or with 1500 (1x) or 3000 (2x) randomly generated genes. See discussion.

**Supplementary Table 1. Related to Figure 1. Quantitative Mass Spectrometry Data for 22q11.2 Genealogical Proteomic Studies.**

**Supplementary Table 2. Related to Figure 2A-D. Bioinformatic Analysis of 22q11.2 Genealogical Proteomes and *Df(16)A*-/+ brain proteomes.**

**Supplementary Table 3. Related to Figure 2E-F. Quantitative Mass Spectrometry Data for *Df(16)A*-/+ brains.**

**Supplementary Table 4. Related to Figure S1. Comparative Bioinformatic Analysis of 22q11.2 Genealogical Proteomes.**

**Supplemental Table 5. Primers Used in these Studies.**

